# Antiviral and Anti-inflammatory Effects of Cannabidiol in HIV/SIV Infection

**DOI:** 10.1101/2025.09.25.678534

**Authors:** AL Ellison, JJ Rosado-Franco, L Mamun, S Knerler, M Daniali, K Mehta, S Williams-McLeod, J Alvarez, SS Joyner, R Vandrey, EM Weerts, PJ Gaskill, MJ Corley, LC Ndhlovu, DW Williams

**Affiliations:** Department of Pharmacology and Chemical Biology, Emory University School of Medicine, Atlanta, GA, USA; Department of Pharmacology and Physiology, Drexel University College of Medicine, Philadelphia, PA, USA; Cellular and Molecular Medicine PhD Program, Johns Hopkins University School of Medicine, Baltimore, MD, USA; Pathobiology PhD Program, Johns Hopkins University School of Medicine, Baltimore, MD, USA; Neuroscience PhD Program, Florida State University, Tallahassee, FL, USA; Department of Psychiatry and Behavioral Sciences, Johns Hopkins University School of Medicine, Baltimore, MD, USA; Department of Medicine, University of California San Diego, San Diego, CA, USA; Department of Medicine, Division of Infectious Diseases, Weill Cornell Medicine, New York, NY, USA

## Abstract

Persistent reservoirs and chronic immune activation are hallmarks of HIV, despite the effectiveness of antiretroviral therapy (ART) in suppressing viral replication. Here, we use rhesus macaques and primary and induced pluripotent stem cell (iPSC)-derived human immune cells to evaluate the virologic and immunologic consequences of cannabidiol (CBD) exposure during HIV/SIV infection. We show that CBD, in the absence of ART, suppresses viral replication and establishment of the viral reservoir to levels comparable with first-line therapies during acute SIV infection of rhesus macaques. This antiviral effect of CBD extended to *in vitro* HIV infection of human macrophages, T cells, and microglia. Immunologically, we observe CBD slowed CD4+ T cell decline and polarization, decreased CD14+CD16+ monocyte expansion, and reduced interferon-inducible cytokine release in rhesus macaques. We identify comparable effects on cytokine production with *in vitro* CBD treatment of human macrophages, T cells, and microglia. Importantly, we find CBD inhibits cytokines only when an immune response is elicited by HIV, suggesting it is not broadly immunosuppressive. Finally, we determine CBD regulates endocannabinoid receptors, modulators, and transporters and inhibits NF-κb and STAT1 activation when mediating its antiviral and anti-inflammatory effects. These findings show beneficial effects of CBD in laboratory models of untreated HIV, thus placebo-controlled clinical trials to evaluate the safety and effectiveness of adjunctive CBD use with ART is warranted.

## Main Text

Viral reservoirs and immune dysfunction are barriers to reducing multimorbidity and developing an effective cure, adversely impacting quality of life for people living with HIV despite the success of antiretroviral therapy (ART) in quelling viral replication and reducing mortality^1,2^. Even with consistent long-term use, ART cannot prevent viral rebound after treatment interruption or restore immune activation to pre-infection levels. Thus, latent virus and chronic immune activation remain pressing challenges in the modern HIV era and contribute to the development of comorbid conditions that are risk factors for poor long-term outcomes, including neurologic disease^3^. There are no existing clinical interventions that suppress residual HIV replication, address latency, and hinder ongoing immune activation despite substantial efforts to purge or “block” the latent reservoir, suppress sustained immune responses, and create next-generation ART^4,5^.

The endocannabinoid system is recognized as an important regulator of many physiological responses, including immune homeostasis, and has emerged as a potential therapeutic target for immune dysfunction^6^. The endocannabinoid system is comprised of lipid ligands, receptors with which they interact, modulators that regulate their synthesis and degradation, and transporters that facilitate their entry into cells^7^. Canonical endocannabinoid receptors (CB) include CB1 and CB2. However, endocannabinoid ligands can bind to additional proteins whose primary functions are involved in other pathways, termed extended endocannabinoid receptors. These extended endocannabinoid receptors include peroxisome proliferator-activated receptor (PPAR)-α, PPAR-γ, transient receptor potential vanilloid (TRPV)-1, TRPV-2, G protein-coupled receptor (GPR)-18, GPR55, GPR119, GPR110/ADGRF1, serotonin 1A receptor (5-HTR1A), and adenosine A2A receptor (ADORA2A)^8^. Cannabidiol (CBD) is a primary constituent of cannabis and elicits downstream immunomodulatory effects by interacting with endocannabinoid receptors. CBD is a non-intoxicating cannabinoid with an excellent safety profile and low abuse liability^9–12^. As such, there is much interest in its clinical promise. The US Food and Drug Administration (FDA) has already approved an oral formulation of purified CBD (Epidiolex©) for treatment of seizures associated with Lennox-Gastaut syndrome, Dravet syndrome, or tuberous sclerosis complex^13^. For these reasons, we aimed to evaluate whether CBD’s immunomodulatory potential could be useful in ameliorating immune dysfunction in the context of HIV infection.

Here, we evaluated the virologic and immunologic consequences of a pure CBD isolate *in vivo* with a rhesus macaque model of acute simian immunodeficiency virus (SIV) infection and *in vitro* following HIV infection of primary and iPSC-derived human immune cells. Using a once-weekly oral administration regimen, we determined that just three doses of CBD decreased plasma and cerebrospinal fluid (CSF) viral replication and the tissue latent reservoir to levels that rivaled suppression afforded by six months of ART treatment. Flow cytometry determined that CBD administration during acute SIV infection maintained peripheral blood and spleen cellular immunity, evidenced by preventing CD4+ T cell decline and polarization into effector subsets and decreasing CD14+CD16+ monocyte expansion. CBD’s *in vivo* immunomodulatory effects extended to decreasing plasma, CSF, and tissue interferon (IFN)-induced cytokines. Our *in vitro* studies with primary human cells determined that CBD’s antiviral and anti-inflammatory capacities occurred in macrophages, T cells, and microglia, though cell-type- and donor-dependent differences occurred. We found that CBD regulated endocannabinoid receptors, modulators, and transporters *in vivo* among multiple brain regions and peripheral organs, as evidenced by scATAC-seq and qRT-PCR. In contrast, endocannabinoid-related lipids remained relatively unchanged in plasma and CSF, suggesting CBD’s effects were not mediated through ligand modulation but rather through receptor signaling. We tested this hypothesis using primary human macrophages and iPSC-derived microglia and found CB2 agonism mirrored the antiviral effects of CBD. Further, *in vitro* infection of primary human cells determined that CBD inhibited activation of nuclear factor kappa B (NF-κb) and signal transducer and activator of transcription 1 (STAT1), transcription factors essential for HIV/SIV transcription and the antiviral immune response. These findings provide a measured optimism regarding the potential clinical utility of CBD for people living with HIV, with implications in viral persistence, cure strategies, and restoring chronic immune activation.

### CBD Decreases SIV Infection *in vivo*

Using an oral administration regimen, we first assessed whether CBD alone could influence SIV infection and the viral reservoir to a similar extent as a current first-line ART regimen. Rhesus macaques were infected with SIV_mac251_ for one week, received CBD once weekly for three weeks, and the study terminated at four weeks post-inoculation (**Figure 1A**). We observed that CBD administration decreased the SIV p27 antigen in both plasma (**Figure 1B**) and CSF (**Figure 1C**) at the study end relative to the point of maximal production. Similarly, plasma (**Figure 1D**) and CSF (**Figure 1E**) SIV_Gag_ RNA were significantly decreased at the terminal time point for SIV-infected animals that received CBD compared to those that did not receive CBD. Surprisingly, SIV_Gag_ RNA was comparable between infected animals that received CBD and those that received ART. CBD also impacted the viral reservoir, as SIV_Gag_ DNA was decreased in tissues (**Figure 1F**) and single-cell isolates (**Figure 1G**) from anatomic sites relevant for SIV/HIV latency compared to the median value from SIV-infected animals that did not receive CBD (**Figure 1F**, black dashed line). Notably, after three weeks of CBD administration, SIV_Gag_ DNA was lower in multiple brain regions, liver, and some lung, kidney, spleen, lymph node, and colon specimens relative to the median value (**Figure 1F**, blue dashed line) from animals that achieved complete viral suppression with ART (**Supplemental Figure 1**). CBD did not affect animal weight, subclinical anemia, or platelet decline characteristic of acute SIV infection^14–16^ (**Supplemental Figure 2**). To our knowledge, this is the first demonstration of the potent antiviral capacity of CBD *in vivo*.

**Figure 1:**
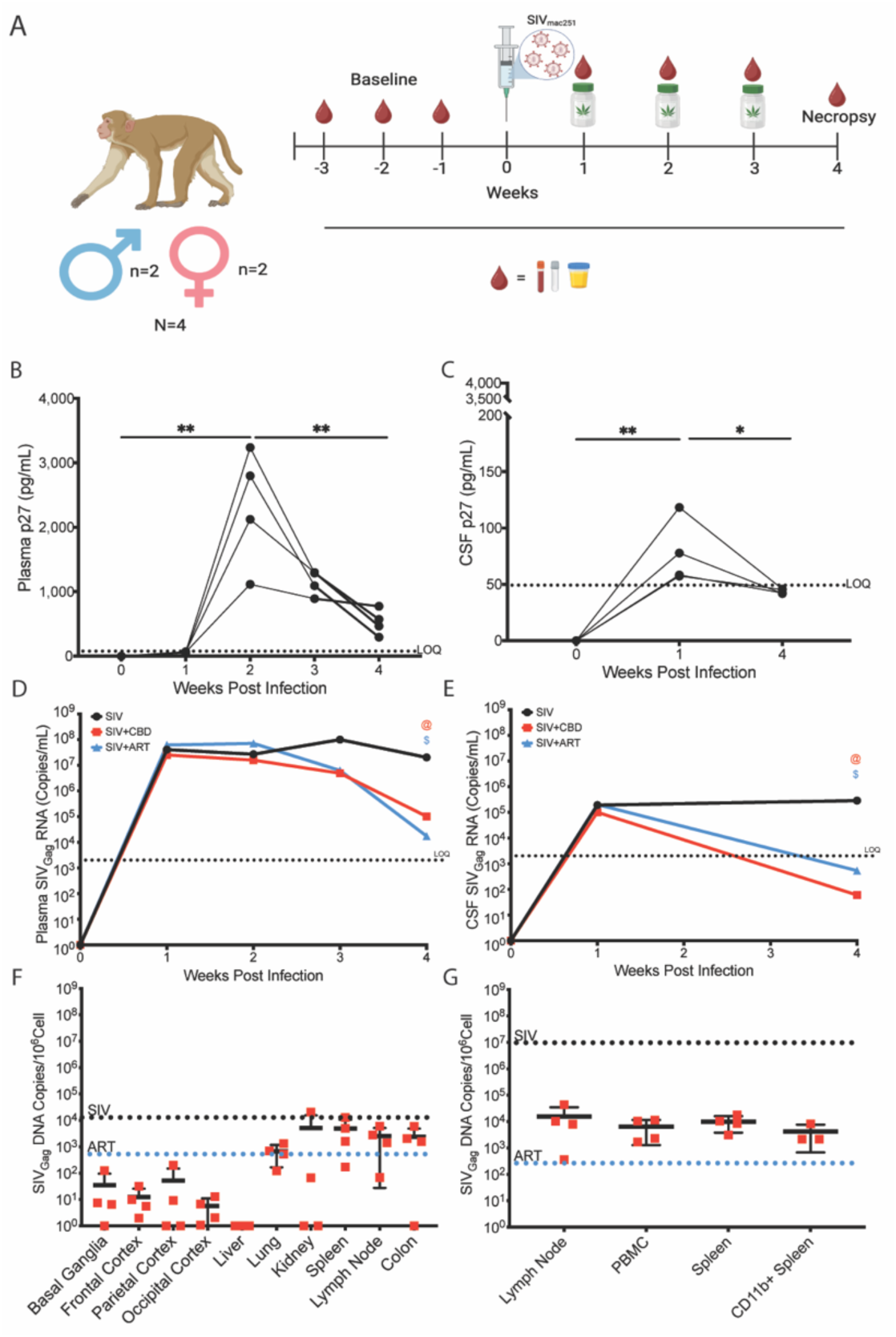
CBD Decreases Viremia and Establishment of the Latent Reservoir During Acute SIV Infection. (A) Schematic of oral CBD administration rhesus macaque paradigm. Four rhesus macaques (n=2 female, n=2 male) received once weekly blood draws prior to study initiation. The macaques were SIV-infected and received once-weekly oral CBD administration with longitudinal sampling of blood, urine, or CSF for three weeks. The study was terminated and necropsy performed in the fourth week. (B-C) Longitudinal SIV p27 capsid antigen determination in (B) plasma and (C) CSF in the SIV-infected, CBD-treated macaques. Data illustrate p27 values at each week post infection where individual macaques (n=4) are indicated by black circles. *p≤0.05. **p≤0.01. Two-tailed, unpaired T test. The dashed line indicates the limit of detection (LOD). (D-E) Longitudinal SIV_Gag_ RNA determination in (D) plasma and (E) CSF from SIV-infected macaques (black), SIV-infected, CBD-treated macaques (red), or SIV-infected, ART-treated macaques (blue). Each macaque group consisted of n=4 animals (n=2 female, n=2 male). Data are represented as median SIV_Gag_ RNA values. @p0.01 for SIV-infected, CBD-treated macaques at week four relative to SIV-infected macaques. $p≤0.01 for SIV-infected, ART-treated relative to SIV-infected macaques. Two-tailed, unpaired T test. The dashed line indicates the limit of detection (LOD). (F-G) SIV_Gag_ DNA determination in (F) tissue punches or (G) single cells isolated from the indicated organs at the terminal time point from SIV-infected, CBD-treated macaques (red). Data are represented as mean±standard deviation. The black dashed line indicates median SIV_Gag_ DNA values from SIV-infected macaques^20,23,25,27,40,47–49^. Blue dashed line indicates median SIV_Gag_ DNA values from SIV-infected, ART-treated macaques^20,23,25,27,40,47–49^.

### CBD is Immunoprotective *in vivo*

We next assessed the immunomodulatory potential of CBD to restore cellular immunity during acute SIV infection. We first confirmed our SIV infection model caused anticipated decreases in the percentage of peripheral blood CD3+ (**Figure 2A, black**) and CD4+ T cells (**Figure 2B, black**), and a resultant increase in CD8+ T cells (**Figure 2C, black**) at 1-week post-infection compared to the baseline time point. Further, we ensured acute SIV infection promoted CD4+ T cell polarization from naïve to effector subsets (**Figure 2D, black**) and increased the frequency of CD14+CD16+ monocytes (**Figure 2F, black**) in peripheral blood, as is well-established^17–20^. We confirmed these expected SIV-induced changes in cellular immunity were not restricted to blood but also occurred in the spleen, a lymphoid organ relevant for HIV/SIV pathogenesis^21^ (**Figure 2G-I, black**). Remarkably, these cellular hallmarks of acute infection did not occur in the blood or spleen of SIV-infected animals that received CBD (**Figure 2A-C and E-I**, **red**).

**Figure 2:**
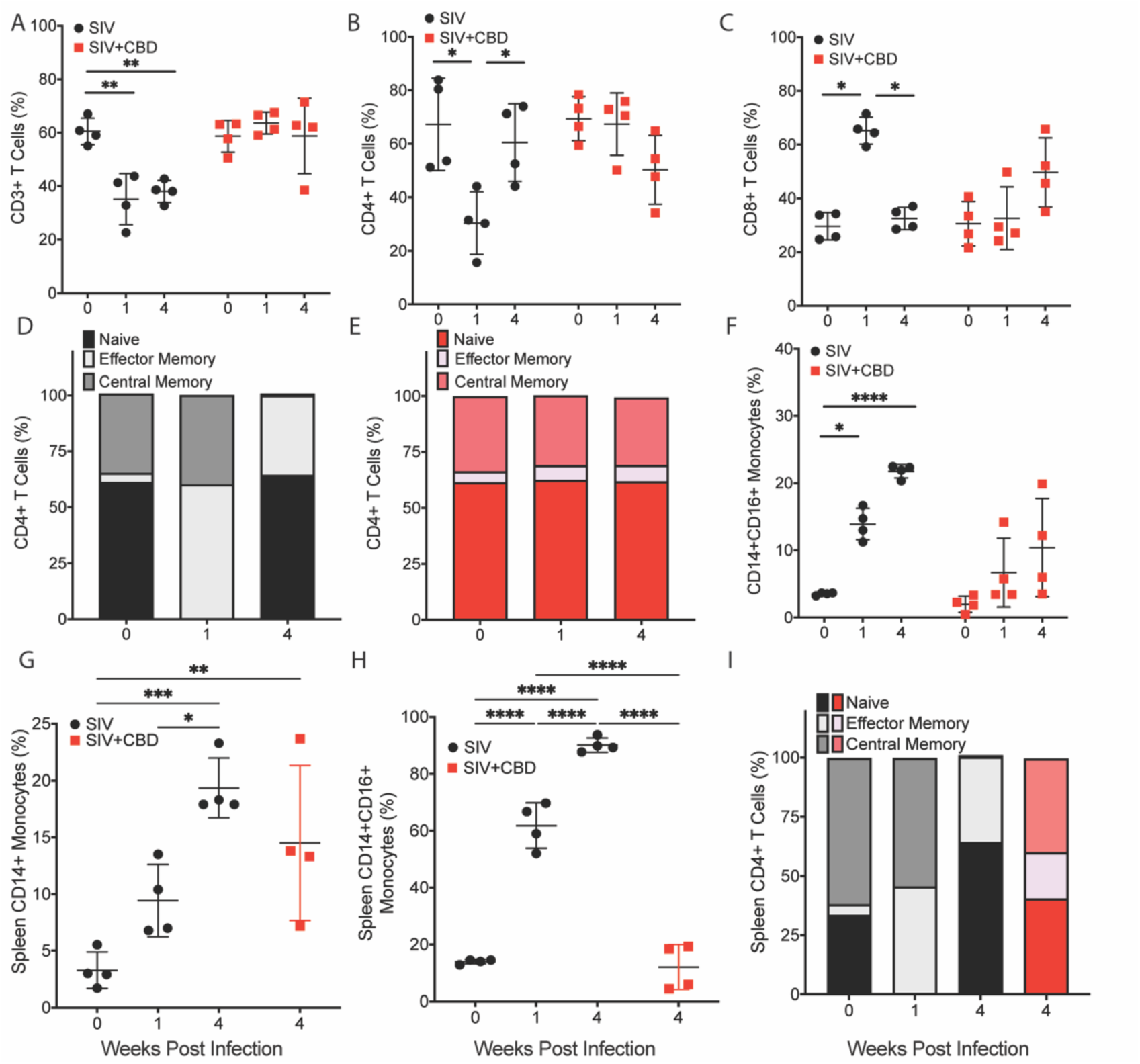
CBD Prevents Peripheral Blood and Spleen Immune Cell Activation During Acute SIV Infection. Percentage of peripheral blood (A) CD3+ T cells, (B) CD4+ T cells, (C) CD8+ T cells, (D-E) CD4+ T cell effector subsets, and (F) CD14+CD16+ monocytes from SIV-infected (black) or SIV-infected, CBD-treated (red) macaques at 0-, 1-, and 4-weeks post-inoculation. Percentage of splenic (G) CD14+ monocytes, (H) CD14+CD16+ monocytes, and (I) CD4+ T cell effector subsets from SIV-infected (black) or SIV-infected, CBD-treated (red) macaques at 0-, 1-, and 4-weeks post-inoculation. Data are represented as mean±standard deviation. *p≤0.05. **p≤0.01. ***p≤0.001. ****p<0.0001. One-way ANOVA with Tukey’s multiple comparisons test.

We then evaluated IFN-inducible cytokines to assess whether CBD could suppress the cytokine storm characteristic of acute viral infection. Plasma interleukin (IL) IL-1β, IL-6, tumor necrosis factor-α (TNF-α), IFN-γ, IFN-γ inducible protein 10 (IP-10), and granulocyte-macrophage colony-stimulating factor (GM-CSF) were significantly decreased at the terminal timepoint relative to that of maximal cytokine production in SIV-infected animals that received CBD (**Figure 3A and 3C; Supplemental Figure 3**). CBD’s effects on IFN-inducible cytokines were more prominent in CSF. All cytokines evaluated were significantly decreased at the terminal timepoint, including IL-8, IL-12p70, IFN-α, IFN-β, and IL-10, which were not vulnerable to modulation in plasma (**Figure 3B and 3D; Supplemental Figure 4**). There was the notable exception of IP-10, which decreased but did not reach statistical significance due to variability in its CSF concentrations among animals (**Supplemental Figure 4K**). In addition to circulating cytokines, we also evaluated select cytokines and IFN-inducible transcription factors in spleen, kidney, liver, mesenteric lymph node, adipose tissue, and heart to determine whether CBD’s immunomodulatory effects extended to diverse organ compartments. C-C motif chemokine ligand 2 (CCL2), TNF-α, IL-12a, and tumor necrosis factor-related apoptosis-inducing ligand (TRAIL) were significantly decreased in an organ-specific manner in SIV-infected animals that received CBD relative to those that received vehicle (**Figure 3E-H**). Interestingly, the IFN transcription factor myxovirus resistance protein A (MxA) remained unchanged with CBD administration (data not shown). Combined, these findings demonstrate that CBD possesses the selective ability to decrease the acute immune response to SIV.

**Figure 3:**
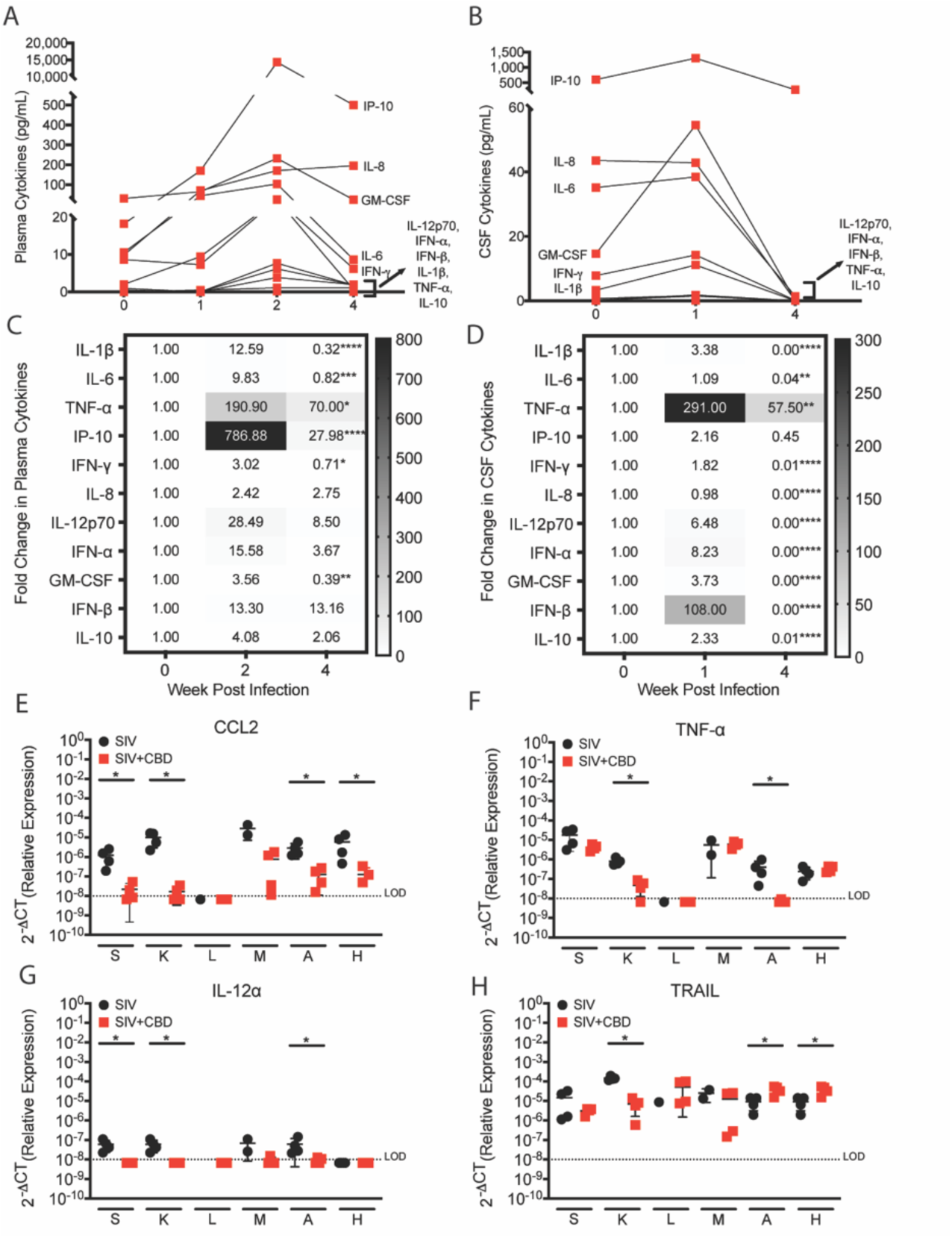
CBD Decreases Interferon-Induced Immune Response During Acute SIV Infection. (A-D) Longitudinal (A, C) plasma and (B, D) CSF cytokine determination in SIV-infected, CBD-treated macaques. (A-B) Data are represented as median cytokine values of n=4 macaques. (C-D) The mean cytokine concentration at week 0 post-infection was determined, set to 1, and the fold change relative to that value was determined for cytokine production at weeks 2 and 4 post-inoculation. Numbers indicate mean fold change relative to week 0 post-inoculation. Data are represented as mean±standard deviation. *p≤0.05. **p≤0.01. ***p≤0.001. ****p<0.0001. Friedman test with Dunn’s multiple comparison’s test. (E-H) Relative gene expression for (E) CCL2, (F) TNF-a, (G) IL-12a, and (H) TRAIL in spleen (s), kidney (k), liver (l), mesenteric lymph node (m), adipose tissue (a), and heart (h) from SIV-infected (black) and SIV-infected, CBD-treated macaques (red). The dashed line indicates the limit of detection (LOD). *p≤0.05. Two-tailed Mann-Whitney test.

### CBD Selectively Modulates Endocannabinoid Tone *in vivo*

We next compared the status of the endocannabinoid system in uninfected, SIV-infected, and SIV-infected animals that received CBD. First, we evaluated CB1 (*cnr1)*, CB2 (*cnr2*), FAAH (*faah)*, NAAA (*naaa)*, PPAR-α (*ppar*α), PPAR-γ (ppar*γ*), TRPV-1 (*trpv1*), TRPV-2 (*trpv2*), GPR18 (*gpr18*), GPR55 (*gpr55*), GPR119 (*gpr119*), GPR110/ADGRF1 (*gpr110*), 5-HTR1A (*htr1a*), and ADORA2A (*a2ar*) gene expression in five brain regions and six organs by qRT-PCR. Surprisingly, SIV infection did not change any of the 14 endocannabinoid receptors, modulators, and transporters in the evaluated organs and brain regions, relative to uninfected macaques (data not shown). However, CBD administration significantly modulated eight genes in peripheral organs (**Supplemental Figure 5A-H**) and brain (**Supplemental Figure 5I-P**) in an organ- and region-specific manner compared to SIV-infected animals that did not receive CBD. Notably, CBD modulated CB1, PPAR-α, PPAR-γ, and TRPV-2 in most evaluated brain regions and organs. Next, we evaluated 33 endocannabinoid-related lipid ligands longitudinally in the plasma and CSF of SIV-infected animals that received CBD. These lipid ligands remained remarkably stable during infection and following CBD administration, where only one ligand was changed in each compartment (**Supplemental Figure 6)**. This is in striking contrast to the robust modulation of endocannabinoid receptors, modulators, and transporters that occurred with CBD.

Finally, we performed single-cell multiomics (scRNA-Seq + scATAC-seq) to more comprehensively evaluate CBD’s effects on the endocannabinoid system. We compared hippocampi from uninfected animals with SIV-infected macaques that received CBD, focusing on endocannabinoid receptors, modulators, and transporters. Consistent with our qPCR findings, CB1 was enriched in SIV-infected animals that received CBD, which was primarily restricted to neuronal populations (**Figure 4**). In contrast, CB2, GPR18, GPR55, GPR119, and TRPV1 were not differentially expressed. In addition to endocannabinoid receptors, CBD administration impacted endocannabinoid modulators responsible for anandamide synthesis (ANA), 2-arachidonoylglycerol (2-AG) degradation, and cytosolic phospholipase A2 (PLA2G4A), which regulates endocannabinoid precursors. We also found selective enrichment of fatty acid binding protein 3 (FABP3) that facilitates endocannabinoid intracellular transport. Collectively, our findings demonstrate that CBD, but not SIV, regulates endocannabinoid tone in a complex fashion in the brain and periphery. These findings raise the question as to whether the endocannabinoid system can be exploited in HIV cure efforts.

**Figure 4.**
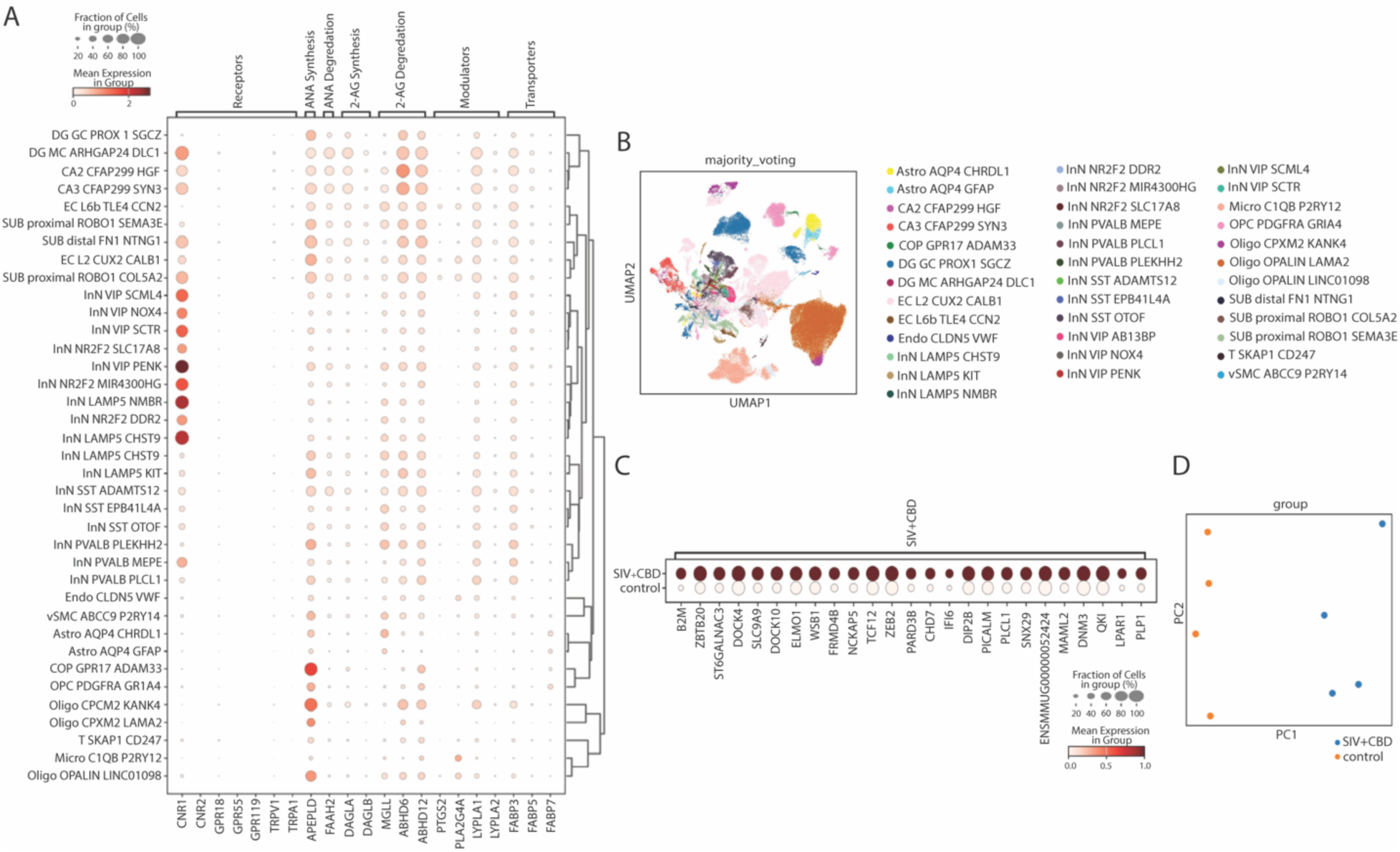
Single-cell transcriptional profiling of hippocampus cell populations. (A) Dot plot showing expression of canonical marker genes across identified hippocampus cell clusters from rhesus macaque specimens annotated based on Franjic et al 2022^50^. Dot size indicates the percentage of cells within a cluster expressing the marker, and color intensity reflects the average expression level. (B) UMAP projection of all single cells, colored by major cell-types identity based on canonical marker gene signatures. (C) Dot plot highlighting the expression of defining marker genes for differential gene expression gene subset, with dot size representing the percentage of cells expressing the marker and color scale indicating mean expression. (D) Principal component analysis (PCA) plot showing separation of rhesus macaque SIV+CBD by experimental group based on pseudo-bulk transcriptional profiles.

### CBD Decreases Macrophage, T Cell, and Microglia HIV Infection *in vitro*

Our finding that CBD decreases SIV infection and inflammatory responses in multiple anatomic sites indicates that the immune cells within these compartments are permissive to cannabinoid-mediated modulation. As lymphoid and myeloid cells are present in all evaluated tissues, and known to express endocannabinoid receptors^6^, we next characterized CBD’s ability to suppress HIV infection of human macrophages, T cells, and microglia (**Figure 5A, Supplemental Figure 7**). All cell types were permissive to HIV infection, with concentrations of the p24 antigen released into culture supernatant varying among donors (**Figure 5B-D**). We found that while CBD decreased concentrations of the HIV p24 antigen released by all cell types, statistically significant changes occurred in macrophages and T cells (**Figure 5B and 5C**) but not microglia due to inter-donor variability (**Figure 5D**).

**Figure 5:**
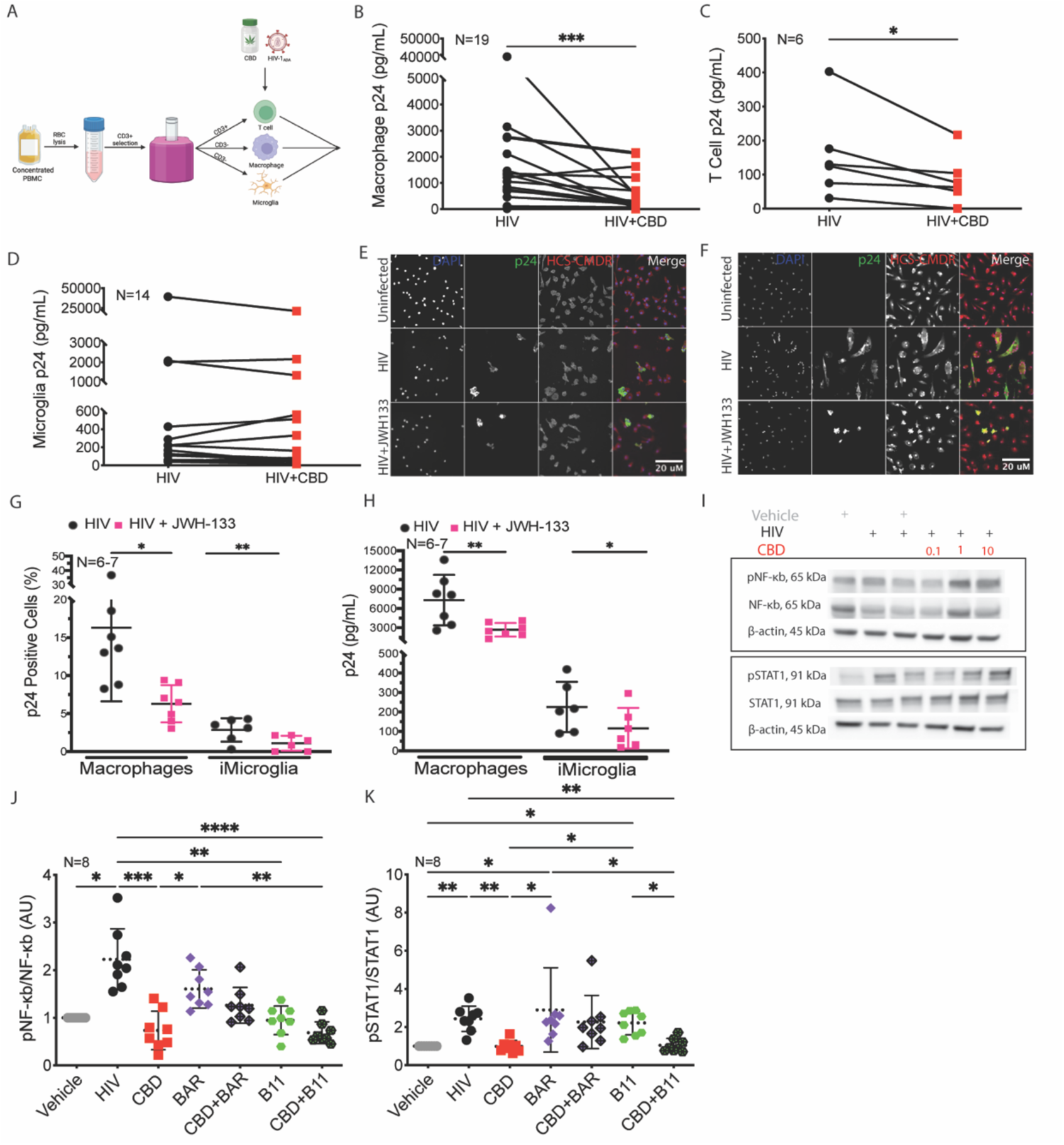
CBD Decreases HIV-Infection and NF-κb and STAT-1 Activation in Human Macrophages, T Cells, and Microglia. (A) Schematic of PBMC isolation and magnetic bead separation to obtain primary human cells for *in vitro* HIV infection and treatments. (B-D) HIV p24 capsid determination in (B) macrophages, (C) T cells, and (D) microglia from HIV-infected (black) and HIV-infected, CBD-treated (red) cells. Mean values for duplicate measures are indicated. N=cells derived from individual human donors. *p≤0.05. ***p≤0.001. Two-tailed Wilcoxon test. (E-F) p24+ cell determination from (E) iPSC-derived microglia and (F) macrophages probed for DAPI, p24 capsid, and HCS CellMask Deep Red (HCS-CMDR). (G) Number of p24 positive cells determined from HIV-infected (black) or HIV-infected, CBD-treated (magenta) macrophages and iPSC-derived microglia (iMicroglia). N=cells derived from individual human donors. Data are represented as mean±standard deviation. *p≤0.05. **p≤0.01. Two-tailed T test. (I) Image from one macrophage donor, representative of eight independent human donors, whose cells were treated with vehicle (grey), HIV (black), or CBD (red), evaluating phosphorylated or total NF-κB or STAT-1. (J-K) Densitometric of (J) phosphorylated/total NF-κB and (K) phosphorylated/total STAT1 from macrophages treated with vehicle (grey, fold change determined, set to 1), HIV (black), CBD (red), Baricitinib (BAR, purple), CBD and Baricitinib (hatched purple), Bay-11 (green), or CBD and Bay-11 (hatched green). N=cells derived from individual human donors. Data are represented as mean±standard deviation. *p≤0.05. **p≤0.01. ***p≤0.001. ****p<0.0001. Kruskal-Wallis test with Dunn’s multiple comparisons test.

These results raise questions regarding the molecular mechanisms by which CBD’s antiviral effects occur. Therefore, we evaluated the contribution of proteins implicated in cannabinoid-mediated immunomodulation. First, we evaluated CB2 using the selective agonist, JWH133, in iPSC-derived microglia and primary human macrophages. We found that CB2 agonism significantly decreased the percentage of cells expressing p24 (**Figure 5E-5G**) and the concentration of p24 antigen released into the supernatant (**Figure 5H**) from HIV infected cultures for both cell types. Next, we performed Western blot to examine CBD’s effects on activation of NF-κb and STAT1, key players in regulating antiviral and innate immune responses. We determined that CBD decreased HIV-induced NF-κb and STAT1 activation in macrophages (**Figure 5I-5K**) and T cells (data not shown). Furthermore, inhibition of NF-κb and STAT1 using Bay 11 and Baricitinib, respectively, showed only partial reduction in NF-κb and STAT1 activation when cells were also treated with CBD. While CBD’s ability to inhibit NF-κb and STAT1 activation was concentration independent, we found that each donor had a unique concentration (between 0.1-10 μM) that maximally inhibited the proteins’ phosphorylation. The 1 μM concentration was most consistent among donors (data not shown) and was subsequently used for remaining downstream experiments. Collectively, these results indicate that CBD’s antiviral effects extend to human immune cells and its mechanistic underpinnings are likely multifactorial, with some effects driven by activation of endocannabinoid receptors, like CB2, and others derived from reduced immune activation, as was the case with NF-κb and STAT1.

### CBD Decreases Macrophage, T Cell, and Microglia Cytokines *in vitro* Only When Induced By HIV

We next probed CBD’s anti-inflammatory capacity in primary human macrophages, microglia, and T cells. Cells were infected with HIV, or remained uninfected as a control, the cells treated with CBD or vehicle, and supernatant was collected to quantify IFN-inducible cytokines (**Supplemental Figure 7**). We found an unanticipated dichotomous immune response to HIV. Infection promoted cytokine secretion in some donors, coined “cytokine inducers”, while it caused no change or even decreased cytokines for the remaining donors, whom we termed “cytokine non-inducers” (**Figure 6A-6C**). There were no distinguishable differences in age, sex, blood type, or ancestry demographics between the donors (data not shown). Further, there was no difference in the concentration of p24 released into the supernatant between “cytokine inducers” and “cytokine non-inducers” (data not shown). Additionally, denotation as a “cytokine inducer” was cytokine specific such that the same individual could be labeled an “inducer” for one cytokine but a “non-inducer” for another, decreasing the likelihood that there were substantial genetic differences between the groups. In fact, only one donor was identified as a “cytokine inducer” for all cytokines across all cell types.

**Figure 6:**
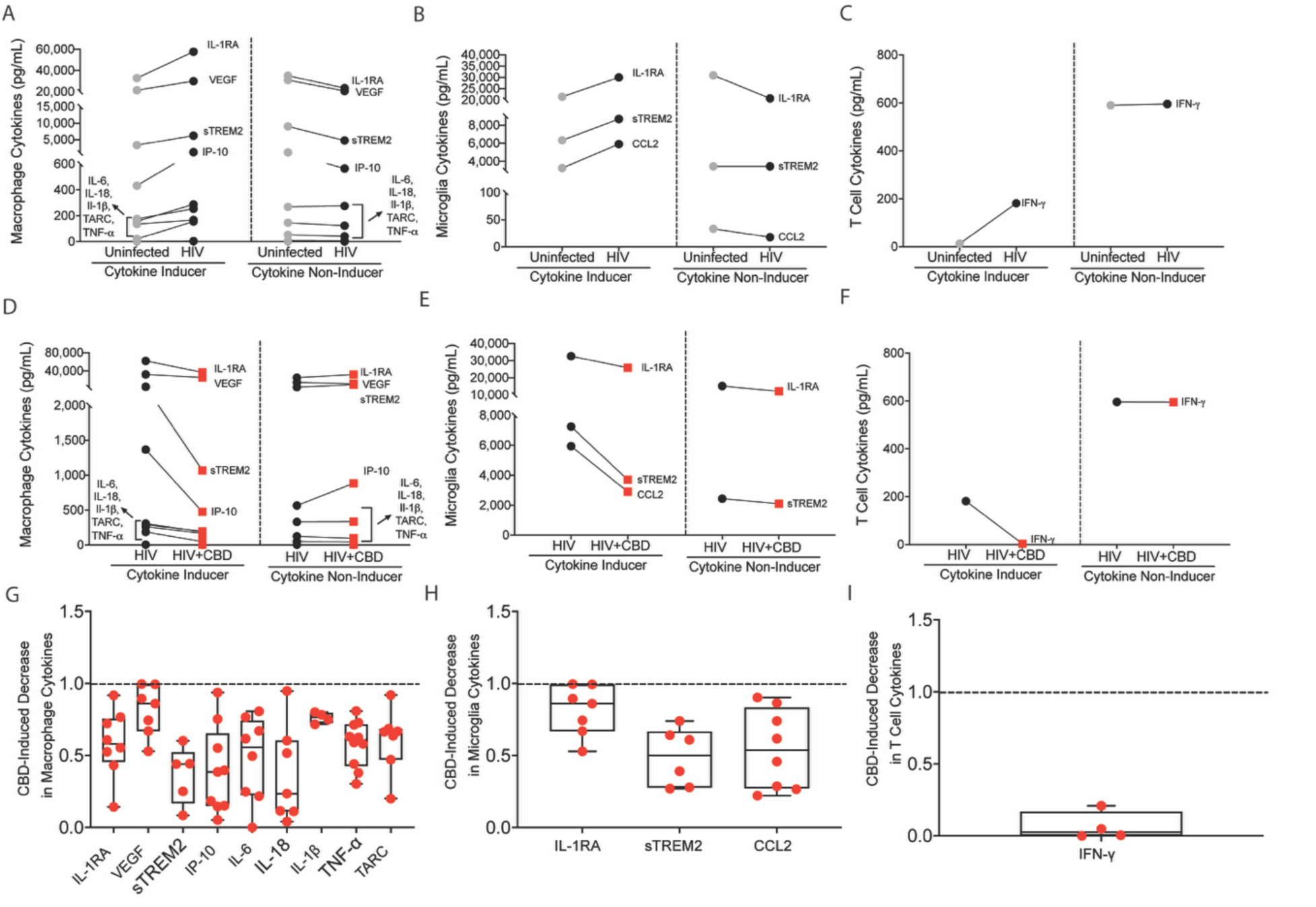
CBD Decreases HIV-Induced Cytokines. (A-C) Cytokine determination in supernatants collected from uninfected (grey) and HIV-infected (black) (A) macrophage, (B) microglia, and (C) T cell cultures. Median values are shown. (D-E) Cytokine determination in supernatants collected from HIV-infected (black) and HIV-infected, CBD-treated (red) (D) macrophage, (E) microglia, and (F) T cell cultures. Median values are shown. (G-I) The average cytokine concentration for HIV-infected cells was calculated, set to 1, and the fold decrease for HIV-infected, CBD-treated (G) macrophages, (H) microglia, and (I) T cells was determined. Represented data are only from the “cytokine inducers” and are depicted as the minimum to maximum value, where each point indicates cells from an individual donor. CBD induced a significant decrease (p≤0.05) for each indicated cytokine, relative to HIV-infected cells, as determined by Two-tailed Wilcoxon test (macrophages and microglia) or Two-tailed T-test (T cells).

We found that this dichotomous cytokine response also occurred following CBD treatment. CBD treatment significantly decreased IFN-inducible cytokines in all cell types (**Figure 6D-6I**). However, these effects were restricted to the “cytokine inducers”. Importantly, CBD had no effect on cytokine production in the “cytokine non-inducer” cells, suggesting it selectively regulates the immune response only in the context of inflammation but does not broadly compromise host immunity. Overall, this data shows that CBD’s anti-inflammatory effects extend to human immune cells, with the added caveat that, in human cells, these effects are selective for heightened inflammation.

## Discussion

Though ART effectively addresses viral replication, there are no clinical interventions for HIV that effectively address the latent reservoir or chronic immune activation, despite substantial efforts. Here, we identify CBD as an antiviral and anti-inflammatory mediator with potential for therapeutic benefit during HIV. To our knowledge, this is the first demonstration of the ability of CBD to reduce early seeding of the viral reservoir, decrease HIV/SIV replication, prevent cellular activation, and suppress IFN-mediated immune responses *in vivo*. We also found striking similarities between human and non-human primates, demonstrating the translational nature of our work. Together, our findings merit cautious optimism regarding the adjunctive utility of CBD during HIV and warrant further investigation in well-controlled clinical trials.

The antiviral capacity of CBD against actively replicating virus and its ability to reduce early seeding of the viral reservoir was unexpected and striking, particularly when compared to ART. There are several well accepted techniques used to measure the viral reservoir, including the Tat/rev Induced Limiting Dilution Assay (TILDA), which assesses the propensity of reactivated CD4^+^ T cells to synthesize virus by measuring tat/rev that is required for intact, functional viral particles^22^. TILDA is more accurate than other measures of the viral reservoir due to its focus on measuring tat/rev, which is required for functional virus and present in only a small portion of nonintact viral particles. Though our measures of SIV_Gag_ DNA overestimate the amount of intact viral DNA, SIV has more intact genomes and less hypermutated proviruses relative to HIV^23^, lending validity to our evaluation of SIV_Gag_ DNA as an appropriate method to estimate early seeding of the stable latent reservoir. Therefore, the overestimation of intact viral particles measured by SIV_Gag_ DNA may be less important and lend itself to being comparable with TILDA measures of the viral reservoir during HIV infection. Additionally, due to the minimum estimate limitations inherent to the quantitative viral outgrowth assay^24^, measures of SIV_Gag_ DNA offer a more accurate indication of seeding during early acute infection^25–27^. For these reasons, SIV_Gag_ DNA was our sole measure of the viral reservoir, though it would be beneficial to integrate additional analyses into future studies. Our findings indicate that CBD reduced SIV_Gag_ DNA in tissues and single cells to levels below those that occurred during untreated acute infection. Additionally, for the first time, we had the notable finding that CBD suppressed SIV_Gag_ DNA in the brain and liver to a greater extent than that of long-term ART treatment. These findings are important and promising for the potential of CBD to decrease the size of what will become the viral reservoir during chronic infection, especially in difficult to treat areas like the brain^28^, where many therapeutic compounds are less effective due to their diminished ability to cross the blood-brain barrier. In addition to these effects in tissues, we found that CBD also limited seeding of the viral reservoir in immune cells, which included CD11b+ myeloid cells and T cells. These changes in the viral reservoir may have been attributed to the ability of CBD to prevent early infection and dysfunction of T cells, as indicated by the maintenance of naïve CD4+ T cells that have a decreased ability to become infected. These findings have major implications for viral eradication and dysfunctional T cell responses that persist despite long-term ART treatment.

The ability of CBD to modulate the immune response induced by HIV/SIV was just as intriguing as its antiviral capacity. Overall, we determined that CBD was immunomodulatory without indiscriminately suppressing all immune function and, at the cellular level, required induction of an immune response before its anti-inflammatory potential was realized. Cytokines in plasma, CSF, and tissue were markedly reduced after CBD exposure, wherein important distinctions occurred between the compartments. Our findings indicate that CBD may be particularly relevant for maintaining neurologic health during HIV infection as all evaluated cytokines were reduced to basal levels in CSF, which did not occur with plasma. Our studies with human cells also highlighted nuanced anti-inflammatory effects as we identified key interdonor differences in the extent to which CBD was effective. We demonstrated a distinction between “cytokine inducers” that mounted an anti-HIV cytokine response and the “cytokine non-inducers” for whom this did not occur. For reasons that remain incompletely understood, CBD selectively decreased cytokine production only in cells obtained from “cytokine inducers”, demonstrating a previously unrecognized selectivity to the immunomodulatory effects of CBD. Importantly, CBD’s selective cytokine inhibition occurred in macrophages, T cells, and microglia suggesting these effects have the potential to impact multiple immune cell types. These findings also demonstrate the importance of evaluating the immunomodulatory potential of cannabinoids in humans, or their cells *in vitro*, as we did not observe a dichotomous effect in the nonhuman primate model despite it being our closest genetic relative. Our findings regarding the *in vitro* ability of CBD to selectively reduce cytokines in “cytokine inducers” has significant implications. If successfully translated to people, it indicates that CBD could be administered to a heterogenous population and facilitate important, life-changing benefits for those exhibiting sustained immune activation without immunologic harm to those who produce little to no inflammatory response.

Few studies have investigated the impact of CBD on the endocannabinoid system during HIV^29^. Our study provides an analysis of endocannabinoid receptors, ligands, modulators, and transporters across multiple brain regions and peripheral organs. Unexpectedly, we found no change in the 14 evaluated endocannabinoid receptors, modulators, and transporters in any tissue between uninfected and SIV-infected animals. In contrast, CBD promoted marked changes in the endocannabinoid system, including receptors, modulators, and transporters, across multiple organ systems. Indeed, we found that CBD modulated multiple receptors that regulate endocannabinoid activity (CB1, PPAR-α, PPAR-γ, GPR55, GPR110/ADGRF1, TRPV1, TRPV2, 5-HTR1A, and ADORA2A) but not endogenous endocannabinoid-related lipid ligands. These data suggest antiviral and anti-inflammatory effects mediated by CBD during HIV infection occur not only through the canonical CB1 endocannabinoid receptor, but also the extended receptors whose primary functions are involved in other pathways. Interestingly, we observed an unexpected increase in CB1 in every organ and brain region evaluated. It will be important to evaluate CBD’s long-term effects on CB1 with chronic administration, as CBD does not bind to CB1 but rather is a negative allosteric modulator and endocannabinoid tone may be downregulated following repeat exposure^7^.

While we are not the first to evaluate CBD in the context of HIV, other studies had substantial differences in experimental design compared to our current work. Notably, some *in vitro* studies used nearly three-fold higher CBD concentrations^30,31^, immortalized cells, or pretreated with agents that changed immune cell activation state^32,33^. Differences in study design also occurred with other *in vivo* studies, namely models that used CBD/Δ^9^-tetrahydrocannabinol (THC) co-administration^34^ or noninfectious transgenic viral protein models^29^. Despite these differences, our work builds upon the findings of others regarding the beneficial premise of CBD in HIV. Additionally, our findings expand existing knowledge by demonstrating that CBD’s antiviral and anti-inflammatory capabilities occur across organ compartments, cell types, and species. Notably, our *in vivo* CBD oral administration regimen has never been used in any HIV model and more accurately parallels pilot studies in people living with HIV, though marked differences in CBD formulation and chemical composition (e.g., full spectrum products, products containing THC and/or other cannabinoids), purity^35^ and dosing^36^ existed in these studies that yielded pharmacokinetic profiles and bioavailability distinct from our work.

It is important to consider a limitation of our study includes our use of an animal model and *in vitro* cell culture of human cells. As such, and similar to the work of many investigators, our findings in their current state cannot be directly extrapolated to any patient population. Though these model systems have valuable similarities and parallels to humans, there are also factors that model systems cannot fully replicate, including differences in aspects of human physiology, genetic composition, and in the case of cell culture, function in isolation from the interconnected nature of a whole-body system. Therefore, any assumption that our findings translate directly to humans is inappropriate. Large-scale, placebo-controlled human clinical trials are imperative and must be conducted before beginning to consider the use of CBD as a potential adjunctive treatment for HIV.

These considerations remain particularly important as the relative ease of accessing the cannabis plant and cannabis/CBD products containing different concentrations of cannabinoids and other chemicals has increased. Legalization and policy changes have impacted this increased accessibility, as well as medical and recreational use. However, it is important to note that the only FDA-approved CBD product is Epidiolex©. Commercially available CBD products, while widely accessible, are largely unregulated and have not been evaluated for purity and safety. There is a lack of standard protocols for cultivation, processing, manufacturing, and testing requirements for cannabis products. This lack of standardization contributes to variation in the content of cannabinoids and other chemicals present in CBD products available in dispensaries, stores, and through illicit markets across the US. Indeed, many commercially available CBD products also contain significant amounts of THC, which can have undesired side effects (intoxication, cognitive, and motor effects). Thus, CBD products used by the public are generally not comparable to the CBD isolate used for our experiments. Furthermore, even though our findings are promising, this study should not be interpreted as stating that CBD should be used as a therapeutic for HIV at this time. Rather, the work presented here provides support for further investigation in the form of phase 2-3 placebo controlled, randomized clinical trials evaluating the safety and efficacy of CBD use during HIV infection. Moreover, additional evaluation of potential drug-drug interactions with ART and other medications is also necessary^7^. ART is already FDA-approved for treatment of HIV and there is a wealth of evidence supporting its efficacy in reducing viral load and preventing HIV-related mortality, including data from several large, long-standing clinical trials. Thus, CBD should not be considered as a replacement for ART, but instead evaluated as a potential adjunctive therapy to supplement ART treatment of HIV in aspects that remain uncontrolled, including the viral reservoir and dysfunctional immune activation.

## Methods

### RESOURCE AVAILABILITY

#### Lead Contact

Further information and requests for resources and reagents should be directed to and will be fulfilled by the lead contact, Dionna Williams.

### MATERIALS AVAILABILITY

We did not generate new unique reagents in this study.

### DATA AND CODE AVAILABILITY

Source data are provided for all experimental results presented in the main manuscript and supplementary information. Source data are provided with this paper. The raw and processed scRNA-Seq and scATAC-seq data in this study have been deposited in the Neuroscience Multi-omic Archive (NeMO) public database with the persistent identifier (nemo:dat-fgwdt8n) at https://assets.nemoarchive.org/dat-fgwdt8n.

### EXPERIMENTAL MODEL AND STUDY PARTICIPANT DETAILS

#### Microbe Strains

##### Viruses and viral inoculum

The SIVmac251 viral stock was produced by the laboratory of Ronald Desrosiers and expanded by infecting rhesus macaque PBMC as described previously^37^. HIV-1_ADA_ was obtained through the NIH HIV Reagent Program, Division of AIDS, NIAID, NIH: Human Immunodeficiency Virus-1 ADA, ARP-416 contributed by Howard Gendelman.

#### *in vivo* Animal Studies

##### Animal details and ethics statement

Animals and procedures in this study were approved by and in compliance with the Johns Hopkins and Emory University Animal Care and Use Committees. Animal handling and euthanasia were conducted as stated under the NIH Guide for the Care and Use of Laboratory Animals and the USDA Animal Welfare Regulation. Animals were included in the study if they were healthy, adult rhesus macaques (*Macaca mulatta*) that were experimentally naïve and negative for the immunoprotective MHC class I alleles *Mamu-A*01*, *Mamu-B*08*, and *Mamu-B*13*. Macaques were pair-housed in the same facility to minimize any immunologic stress caused by being single-housed and were given *ad libitum* access to standard monkey chow (Teklad Diet 2018; Indianapolis, Indiana, USA) and water. Rhesus macaque cohorts included n=2 per biological sex, n=4 total in this study, for each cohort to detect a 55% change in a dataset that demonstrates a standard deviation of 0.33 (power of 80%) and significance level of 0.05 (R statistical software).

##### Study design and sample collection

The rhesus macaque (*Macaca mulatta*) model used in this study included four cohorts, where each group was comprised of n=4 animals. The first cohort included uninfected animals who received a 50 mL vehicle inoculum. The second and third cohorts were comprised of SIV-infected rhesus macaques and SIV-infected, ART-treated rhesus macaques, respectively. Cohorts two (SIV) and three (SIV/ART) were inoculated intravenously using SIVmac251 as previously described^37^, with the animals in cohort two being euthanized approximately 40 days post-inoculation. For cohort three, a subcutaneous dose of 2.5 mg/kg dolutegravir (DTG, Viiv, London, England, UK), 20 mg/kg tenofovir disoproxil fumarate (TFV, Gilead, Foster City, California, USA), and 40 mg/kg emtricitabine (FTC, Gilead, Foster City, California, USA) was administered once daily for six months beginning day 42 post-infection; euthanasia occurred at approximately 200 days post-inoculation. Blood and CSF sampling were performed weekly before inoculation, weekly during acute infection, and monthly after ART initiation for cohorts two and three. The fourth cohort of animals was comprised rhesus macaques inoculated intravenously with SIVmac251 in the same manner as cohorts two and three. Additionally, beginning one week post-inoculation, animals in the fourth cohort (SIV/CBD) were administered oral escalating weekly doses of CBD (10mg/kg, 20mg/kg, and 40mg/kg) procured from the Johns Hopkins Behavioral Pharmacology Research Unit Pharmacy and prepared as previously described^38^. Briefly, CBD doses were weighed (g/kg) and thoroughly mixed in USP-grade peanut oil (Sigma Aldrich), subsequently mixed with peanut butter, and applied to graham crackers in sandwich form for administration. Consumption of CBD-peanut butter graham cracker sandwiches was observed and verified. For group four animals, sampling of blood and CSF was performed weekly: before inoculation, at the peak of infection, and weekly during CBD administration; animals were euthanized four weeks post-inoculation (**Figure 1A**). Animal weights were ∼6 kg and maintained throughout the study for all animals (**Supplemental Figure 2A**; data not shown). For euthanasia, all animals were sedated using ketamine and administered an overdose of sodium pentobarbital, according to the American Veterinary Medical Association guidelines. Upon euthanasia at ∼8 AM, tissue harvest was performed in the same manner for all animals and completed by 12 PM. Briefly, CSF and peripheral and chest blood were collected, followed by perfusion using 1X phosphate buffered saline. Tissues were subsequently collected, including brain, colon, heart, kidney, liver, lung, lymph nodes, spleen, and visceral adipose tissue (omental). Brain was further dissected by region, including basal ganglia, frontal cortex, occipital cortex, and parietal cortex. Additionally, PBMC were isolated from blood and single cell isolates obtained from lymph nodes and spleen as previously described^39,40^. Experimental controls for this study were determined in three ways. First, uninfected animals served as a control for all experimental groups. Second, the SIV cohort served as a control relative to the SIV/ART and SIV/CBD cohorts for the purpose of determining ART and CBD efficacy, in which case each animal was considered an experimental unit. Third, longitudinal measures were obtained for each rhesus macaque in all cohorts so that outcomes could be compared between different times in the study, in which case the experimental unit was one animal over a period of time.

#### Primary Cell Culture

##### Chemical compounds for cell culture

The following compounds were prepared in their respective diluents: CBD in methanol (Cayman Chemical Company ISO60156; Ann Arbor, MI, USA), Bay 11 (Cell Signaling Technology 78679; Danvers, Massachusetts, USA) in ethanol, Baricitinib (AmBeed 1187594-09-7; Buffalo Grove, Illinois, USA) in DMSO, and JWH-133 in DMSO vehicle (Tocris 1343; Minneapolis, Minnesota, USA). CBD was added to cells for a final concentration of 0.1 μM, 1 μM, or 10 μM in cell culture media. Bay 11 or Baricitinib was added at a final concentration of 10 μM in cell culture media. JWH-1333 was added at a final concentration of 1 μM in cell culture media.

##### Human peripheral blood mononuclear cell isolation

Leukocyte packs containing concentrated PBMC were purchased from BioIVT (Westbury, New York, USA). Contents were washed with sterile Dulbecco’s Phosphate Buffer Saline (DPBS, Gibco 14190235; Waltham, MA, USA) and red blood cells lysed with 1X lysis buffer (10X buffer comprised of 1.5 M ammonium chloride [Sigma Aldrich A9434; St. Louis, MO, USA], 0.01 M sodium bicarbonate [Sigma Aldrich S5761], 0.01 M EDTA disodium [Sigma Aldrich E5134]). Cells were resuspended in EasySep buffer (StemCell Technologies 20144; Vancouver, British Columbia, Canada) and CD3+ magnetic bead isolation performed according to the manufacturer’s protocol (StemCell Technologies 17851). CD3+ cells were cultured according to the T cell protocol described below, and CD3-flowthrough was cultured according to either the macrophage or microglia protocol below. The age of human donors whose blood was used to obtain cells in this study is 44±13 years. Sex distribution of human donors whose blood was used to obtain cells in this study is N=61 individual donors, where 51% of the donors were assigned female at birth.

##### Macrophage cell culture and HIV infection

CD3- cells were cultured adherently in 10 cm dishes (Fisher Scientific 12567650) in macrophage media (DMEM [Life Technologies 11965118; Carlsbad, California, USA], 10% FBS [R&D Systems S11150; MI, Minnesota, USA], 5% human serum [Gemini GBP-100-512; West Sacramento, CA, USA], 1% HEPES [Quality Biological 118089721; Gaithersburg, MD, USA], 1% penicillin/streptomycin [Gibco 15140122)] 1% L-glutamine [ThermoFisher 25030081], 1% M-CSF [R&D Systems 216-MC-100/CF]), at 37°C, 5% CO_2_ for a total of six days, where fresh media was added at day 3. On day six of culture cells were assessed for differentiation, and if elongated/spindle morphology was apparent, they were considered fully differentiated and used in experiments. Each 10 cm dish of cells was considered one unit. Cells were inoculated with HIV-1_ADA_ virus (5 ng/mL) on day six of culture. Twenty-four hours later, on day seven of culture, the media with the HIV-1_ADA_ inoculum was replaced with fresh macrophage media. Cells were subsequently treated daily with CBD from days 7-11. Select plates also underwent a 1-hour preincubation with Bay 11 or Baricitinib. Cell supernatants were collected on days 8-11, and cells were collected for protein isolation on day 11 as described below. All samples were stored at -80°C. All macrophage experiments were repeated in at least n=9 independent experiments, with cells from a single donor constituting one experiment.

##### Microglia cell culture and HIV infection

CD3- cells were cultured adherently in 10 cm dishes Fisher Scientific 12567650) in microglia media (DMEM [Life Technologies 11965118], 10% FBS [R&D Systems S11150], 5% human serum [Gemini GBP-100-512], 1% HEPES [Quality Biological 118089721], 1% penicillin/streptomycin [Gibco 15140122], 1% L-glutamine [ThermoFisher 25030081], GM-CSF [10ng/mL; PeproTech 30023; Waltham, MA], IL-34 [100ng/mL; PeproTech 20034]) at 37°C, 5% CO_2_. On day 14 of culture, cells were assessed for differentiation and if elongated/spindle morphology was apparent, were considered fully differentiated and used in experiments. Each 10 cm dish of cells was considered one unit. Cells were inoculated with HIV-1_ADA_ virus (10 ng/mL) on day 14 of culture. Twenty-four hours later, on day 15 of culture, the media with the HIV-1_ADA_ inoculum was replaced with fresh microglia media. Cells were treated daily with CBD between days 15-19. Select tubes also underwent a 1-hour preincubation with Bay11 or Baricitinib. Cell supernatants were collected on days 16-19 and cells were collected for protein on day 19 as described below. All microglia experiments were repeated in at least n=14 independent experiments, with cells from a single donor constituting one experiment.

##### T cell culture and HIV infection

CD3+ cells were cultured in Greiner tubes (Greiner Bio-One 07000212; Monroe, North Carolina, USA) in T cell activation media (RPMI [Gibco 11875119], 10% FBS [R&D Systems S11150], 5% human serum [Gemini GBP-100-512], 1% penicillin/streptomycin [Gibco 15140122], IL-2 [10ng/ml; StemCell Technologies 78220.1], PHA-L [5 mg/ml; eBioscience 00497793; Waltham, Massachusetts, USA]) at 37°C, 5% CO_2_. Each tube of cells was considered one unit. On day two of culture, media was replaced with T cell maintenance media (T cell activation media without PHA and IL-2) and HIV-1_ADA_ virus (5ng/mL) was added for 24 hours. Twenty-four hours later, on day three of culture, the media with the HIV-1_ADA_ inoculum was replaced with fresh T cell media. Cells were treated daily with CBD between days 3-7. Select tubes also underwent a 1-hour preincubation with Bay11 or Baricitinib. Cell supernatants were collected on days 4-7, and cells were collected for protein on day 7 as described below. All T cell experiments were repeated in at least n=6 independent experiments, with cells from a single donor constituting one experiment.

##### iPSC-Derived Microglia Differentiation

Common Myeloid Progenitor cells derived from Induced Pluripotent Stem Cells (iPSCs) were obtained from the Children’s Hospital of Philadelphia. These cells were differentiated into iPSC-derived microglia (iMG) over 11 days in RPMI media supplemented with M-CSF (25 ng/ml, PeproTech, 300-25-10UG, Cranbury, New Jersey, USA), TGF-β (50 ng/ml, PeproTech, 100-21-10UG, Cranbury, New Jersey, USA), and IL-34 (100 ng/mL, PeproTech, 200-34-10UG, Cranbury, New Jersey, USA). iMG cells (n=6) were derived from four independent donors.

##### Experimental groups for primary cell culture

Experimental groups for macrophage experiments were designated as follows, with the number of plates per group for each individual experiment/donor indicated in parenthesis: uninfected, untreated (U, n=2), HIV-infected, untreated (HIV, n=2), HIV-infected, MeOH vehicle treated (HIV+MeOH, n=1), HIV-infected, EtOH vehicle treated (HIV+EtOH, n=1), HIV-infected, DMSO vehicle treated (HIV+DMSO, n=1), HIV-infected, Bay11 treated (HIV+Bay11, n=1), HIV-infected, Baricitinib treated (HIV+BAR, n=1), HIV-infected, EtOH vehicle and CBD treated (HIV+EtOH+CBD, n=1), HIV-infected, DMSO vehicle and CBD treated (HIV+DMSO+CBD, n=1), HIV-infected, CBD and Bay11 treated (HIV+CBD+Bay11, n=1), HIV-infected, CBD and Baricitinib treated (HIV+CBD+BAR, n=1), HIV-infected, 0.1 μM CBD treated (HIV+0.1 μM CBD, n=1), HIV-infected, 1 μM CBD treated (HIV+1 μM CBD, n=1), HIV-infected, 10 μM CBD treated (HIV+10 μM CBD, n=1). Experimental groups for T cells and microglia were identical to macrophages, with the caveat that only 1 μM CBD treatments were used, as it was identified as the concentration that yielded the most consistent effects among donors in macrophages. Groups with HIV infection alone were compared to uninfected, untreated samples to confirm infection. Subsequently, groups infected with HIV and treated with CBD or an immune inhibitor were compared to their respective vehicle controls to ensure any potential treatment effects were not simply due to the vehicle.

### METHOD DETAILS

#### Complete Blood Count Panel

Rhesus macaque blood was collected in citrate dextrose solution (ACD; Millipore Sigma C3821; Burlington, Massachusetts, USA) and a complete blood count performed using a ProCyte Dx Hematology Analyzer (IDEXX Laboratories; Westbrook, Maine, USA).

#### SIV_Gag_ RNA Determination

SIV_Gag_ RNA was measured by RT-qPCR with a sensitivity of 40 copies/mL in collaboration with the Emory Center for AIDS Research as previously described^41,42^.

#### SIV_Gag_ DNA Determination

SIV_Gag_ DNA was measured by RT-qPCR with a sensitivity of one copy per 850,000 cells within tissues and cells in collaboration with the Emory Center for AIDS Research, as previously described^41,42^.

#### SIV p27 Quantification

Plasma and CSF SIV p27 protein was quantified with an SIV p27 antigen capture assay (ABL Inc 5436; Rockville, Maryland, USA) according to the manufacturer’s protocol with a limit of detection of 50 pg/mL.

#### HIV p24 Quantification

HIV p24 capsid antigen in cell culture supernatants was quantified with an Alliance HIV-1 ELISA Kit (Perkin Elmer NEK050B001KT; Shelton, Connecticut, USA) according to the manufacturer’s protocol with a limit of detection of 6.25 pg/mL.

#### Whole Blood Flow Cytometry

Flow cytometry was performed on whole blood samples from rhesus macaques to quantify T cell and monocyte subsets, as previously described^40^. At least 500,000 singlet events were acquired using a BD LSRFortessa cytometer and Diva software version 9 or BD FACS Lyric and FACS Suite software on the Windows 10 platform (BD Biosciences; Franklin Lakes, New Jersey, USA). Data were analyzed using FlowJo software version 11 (FlowJo; Ashland, Oregon, USA) and gating strategy performed as detailed in **Supplemental Figure 8**.

#### Cytokine Quantification

Cytokines were quantified with the BioLegend LEGENDplex Human Anti-Virus Response Panel V02 with Filter Plate (BioLegend 741269; San Diego, California, USA), Human Macrophage/Microglia Panel with Filter Plate (BioLegend 740502), and the Human Neuroinflammation Panel 1 with Filter Plate (BioLegend 740795) according to the manufacturer’s protocol as previously described^43^.

#### Gene Expression Quantification

RNA was isolated, cDNA synthesized, and qRT-CPR performed in duplicate using TaqMan Fast Universal PCR Master Mix (2x), No AmpErase UNG (ThermoFisher 4367846), and TaqMan Gene Expression Assay Mix according to our established protocol^8^.

#### Endocannabinoid Quantification

Endocannabinoids were quantified in collaboration with the Emory Integrated Metabolomics and Lipidomics Core using an external calibration curve and analytical grade standards. Calibration curves were prepared over the linear range of 0.1-10 nM and the equation of the slope was used for quantification of the corresponding lipid I plasma. Endocannabinoids were isolated using a Biotage Extrahera, resolved using an Infinity II/6495c LC/MS system, and analyzed by Agilent 6495c mass spectrometer operated in negative and positive ion modes according to an established protocol^44^.

#### Western Blot

Macrophages, T cells, or microglia were lysed with RIPA buffer (Cell Signaling Technology 9806S) supplemented with protease/phosphatase inhibitor (Cell Signaling Technology 5872S; Danvers, Massachusetts, USA). Total protein concentration was determined by Bradford Assay (Bio-Rad 5000006) following the manufacturer’s protocol. Forty μg of protein was mixed with loading buffer (2x Laemmli loading buffer/dye (Bio-Rad 1610737) and 2-mercaptoethanol (Sigma Aldrich M6250), electrophoresed on a 4– 15% polyacrylamide gel (Bio-Rad 4561084; Hercules, California, USA), transferred to nitrocellulose membranes (Amersham Biosciences 10600001; Woburn, MA, USA), and proteins of interest evaluated by Western blot according to our established protocol^43,45^.

#### Single-Nuclei Isolation, Library Preparation, and Sequencing

Single nuclei were isolated from ∼30 mg fresh-frozen rhesus macaque hippocampus brain tissue using a published method^46^. Tissue was minced on dry ice, homogenized in nuclei lysis buffer (MilliporeSigma L9286) with a Dounce homogenizer, filtered (70 µm), and centrifuged (600 g, 5 min, 4°C). The pellet was resuspended in 1X hypotonic buffer (10x Genomics, Pleasanton, CA), layered over 30%/40% Iodixanol gradients (10x Genomics), and centrifuged (4,000 g, 30 min, 4°C). The nuclei band was washed, pelleted (500 g, 5 min), and resuspended in 1X nuclei buffer (10x Genomics). Nuclei with >70% DAPI positivity (Countess II, Thermo Fisher were used for downstream processing. For the 10x Multiome ATAC+GEX data generation, nuclei were transposed (10x Genomics), loaded on Chromium Next GEM Chip J, and subjected to GEM generation, barcoding, cleanup, and pre-amplification. ATAC libraries were prepared from 45 µL of pre-amp product; cDNA from 12-cycle amplification was used for 3′ GEX library prep (10x Genomics User Guide CG000338). Libraries were quantified (Qubit 2.0), quality-checked (Agilent TapeStation), and sequenced on an Illumina NovaSeq 6000. GEX libraries: 28 bp Read1, 8 bp i7, 91 bp Read2; ATAC libraries: 50 bp Read1N, 8 bp i7, 24 bp i5, 49 bp Read2N. Fastq files were generated with bcl2fastq. A *Macaca mulatta* cellranger-arc reference was built using Ensembl Mmul_10 (release 104) annotation and genome assembly.

#### Immunofluorescence Staining and High-Content Imaging

At day five post-infection, macrophages and iMG were fixed with 4% paraformaldehyde (Thermo Fisher Scientific, J19943K2) for 10 minutes at room temperature, permeabilized, and incubated with blocking buffer (1% BSA (Fisher Scientific, BP1600100), 0.1% Tween 20 (Fisher Scientific, BP337-500), and 22.52 mg/mL glycine (Fisher Scientific, BP381-1) in 1X PBS) for 30 minutes. Cells were incubated overnight at 4°C with anti-p24-Gag primary antibody (ATCC, 4121, Manassas, Virginia, USA). Following primary antibody incubation and a complete wash with PBS, cells were treated with Alexa Fluor 488-conjugated secondary (Thermo Fisher Scientific, A31620) antibody diluted in blocking buffer for 1 hour at room temperature. Following a PBS wash, all cells were counterstained with DAPI (Thermo Fisher Scientific, D1306) and HCS CellMask Deep Red (Thermo Fisher Scientific, H32721) for 10 minutes and preserved in 1X PBS. High-content imaging was performed using the CX7 (Thermo Fisher Scientific) platform, capturing 20 fields per well at 20X magnification.

## Acknowledgements

This research was funded by the National Institutes of Health under award numbers F31 DA058562 (ALE), R01 DA052859 (DWW), U01 DA058527 (DWW, CM, LN). This work was supported, in part, by the Center for AIDS Research at Emory University P30 AI050409 and T32AI157855. This research is supported, in part, by the Emory National Primate Research Center National Primate Research Center Grant No. ORIP/OD P51OD011132. The Emory National Primate Research Center is supported by the National Institutes of Health, Office of Research Infrastructure Programs/OD [P51OD011132]. This research is supported, in part, by the Emory National Primate Research Center Grant No. U42 PDP11023, National Institutes of Health’s Office of the Director, Office of Research Infrastructure Programs. Data presented in this study were produced as part of the Single Cell Opioid Response in the Context of HIV consortium (SCORCH: RRID:SCR_022600). Publicly accessible data is available at NeMO Archive (RRID:SCR_002001) under identifier nemo:dat-fgwdt8n (https://assets.nemoarchive.org/dat-fgwdt8n). The content is solely the responsibility of the authors and does not necessarily represent the official views of the National Institutes of Health. Images in figures were created using Biorender.

## Author Contributions

DWW conceived the project, designed the experiments, and supervised the study. ALE, JJRF, LM, SK, MD, KM, MJC, SWM, JS, MD, and CM executed experiments. ALE, JJRF, LM, SK, MD, KM, MJC, SWM, MD, and DWW performed experimental analysis. ALE, MJC, LCN, and DWW wrote and edited the manuscript. LM, JJRF, MD, and DWW prepared the figures. RV, EW, and DWW designed the CBD oral administration methodology. RV, EW, PJG, MJC, LCN, LM, MD, and DWW interpreted the data. All authors read, edited, and approved the final manuscript.

## Materials & Correspondence

Correspondence to Dionna W. Williams.

## Statistical Information

Rhesus macaque blood, plasma, CSF, basal ganglia, frontal cortex, occipital cortex, parietal cortex, adipose, colon, heart, kidney, liver, lung, lymph node, and spleen were collected from four (n=2 male, n=2 female) independent rhesus macaques per cohort. At least n=8 (macrophage), n=6 (T cell), or n=6 (microglia) independent experiments were performed for all *in vitro* experiments. Details regarding the number of independent experiments performed are included in all figure legends. All statistical analyses were performed on data from at least four independent animals per macaque cohort or six independent donors for *in vitro* experiments. Data for Figures 1B-E, 3A-D, 6A-F and Supplemental Figure 1 are graphically represented as median values. Data for Figures 1F-G, 2A-C, 2F-H, 3E-H, 5G-H, 5J-K and Supplemental Figures 2 and 5 are graphically represented as mean ± standard deviation. Data for Figure 2D-E, 2I are graphically represented as stacked bars depicting the mean frequencies of each cell subset for n=4 animals per group. Data for Figures 6G-I are represented demonstrating the minimum to maximum value for each individual donor. Remaining figures are graphically represented as individual points for each animal or individual donor. Statistical analyses were performed using Prism software version 10 (GraphPad Software, Inc.; San Diego, California, USA). A two-tailed, T test was performed on the data in Figures 1, 5G-H, and 6I. A one-way ANOVA with Tukey’s multiple comparisons test was performed on the data in Figure 2. A two-tailed Mann-Whitney test was performed on the data in Figures 3, and 5. A two-tailed Wilcoxon test was performed on the data in Figures 5B-D and 6G-H. A Kruskal-Wallis test with Dunn’s multiple comparisons test was performed on the data in Figures 5J-K. A Friedman test with Dunn’s multiple comparisons test was performed on the data in Supplemental Figures 2-4. In rhesus macaque experiments the week 0 timepoint served as a baseline and the week 1 timepoint served as the SIV only timepoint, as samples were taken prior to SIVmac251 inoculation and CBD administration, respectively. Depending on the comparisons being made, each of these timepoints were used as a reference group for multiple comparisons analyses when a one-way ANOVA test was performed. *p≤0.05. **p≤0.01. ***p≤0.001. ****p<0.0001.

## Extended Data

**Supplemental Figure 1:**
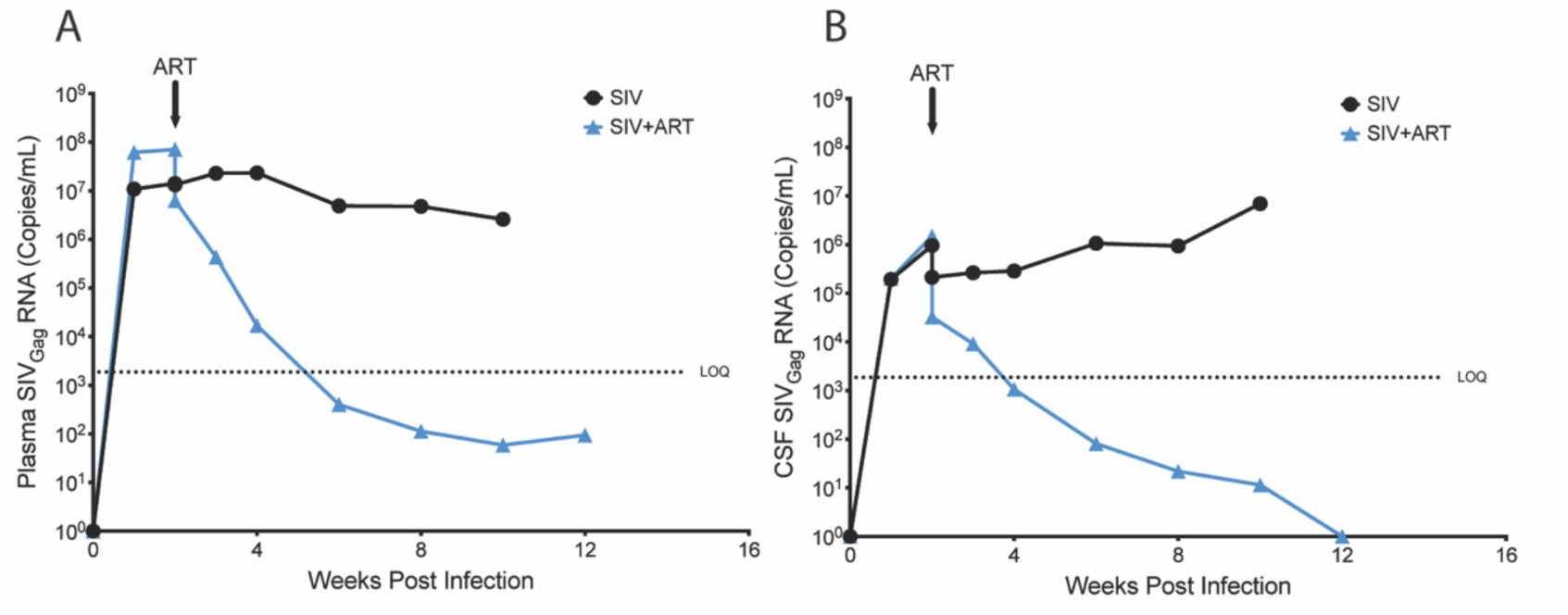
ART Decreases SIV-infection. Longitudinal (A) Plasma and (CSF) SIV_Gag_ RNA determination for SIV-infected (black) and SIV-infected, ART-treated (blue) macaques. Median values for n=4 animals per group are shown. The arrow indicates the time of ART initiation. The dashed line indicates the limit of detection (LOD).

**Supplemental Figure 2:**
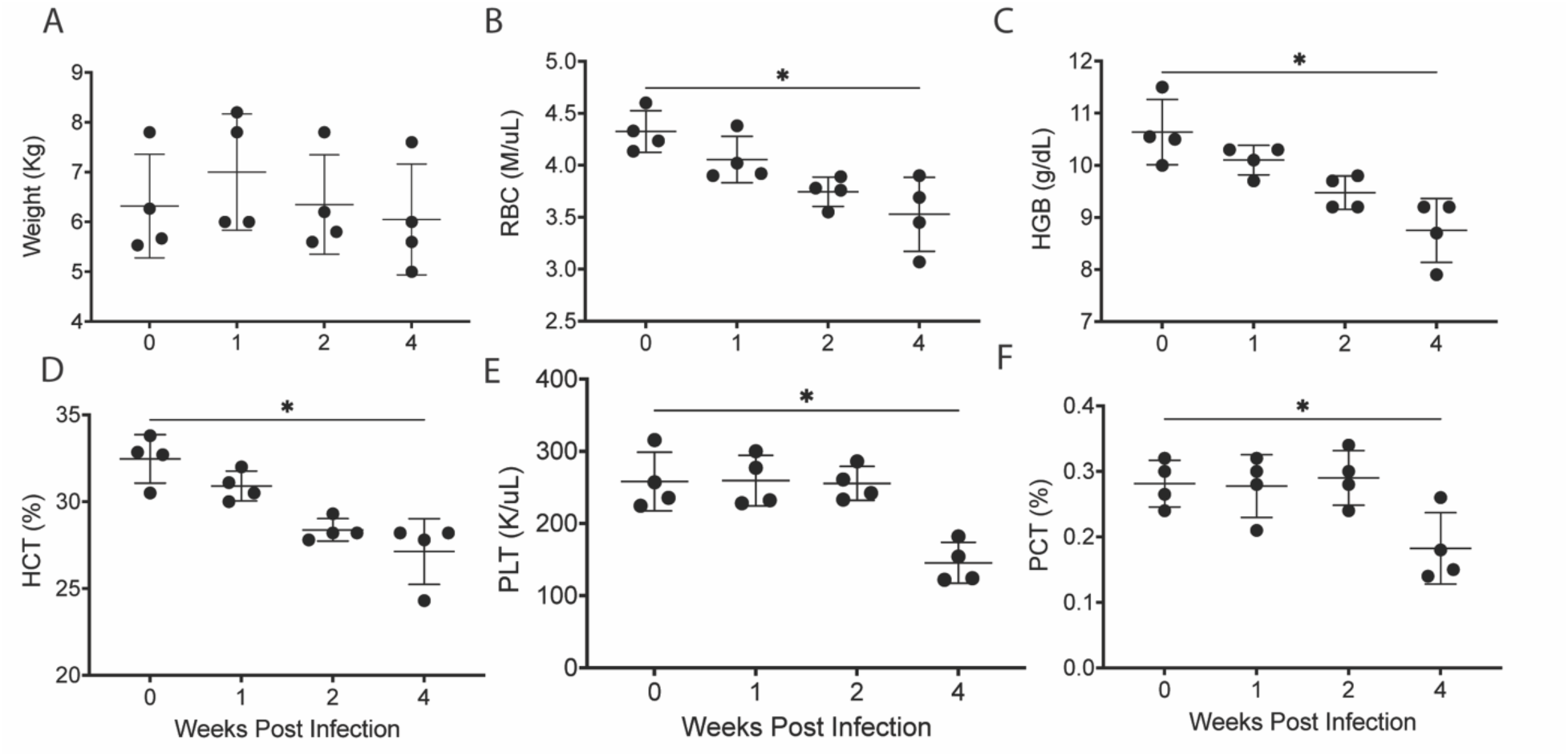
CBD Does Not Change Macaque Weight or Reverse Subclinical Signs of Acute SIV Infection. Longitudinal (A) weight, (B) red blood cell (RBC) count, (C) hemoglobin (HGB), hematocrit (HCT), (E) platelet count (PLT), and (F) plateletcrit (PCT) determination for SIV-infected, CBD-treated macaques. Data are represented as mean±standard deviation. *p≤0.05. Friedman test with Dunn’s multiple comparisons test.

**Supplemental Figure 3:**
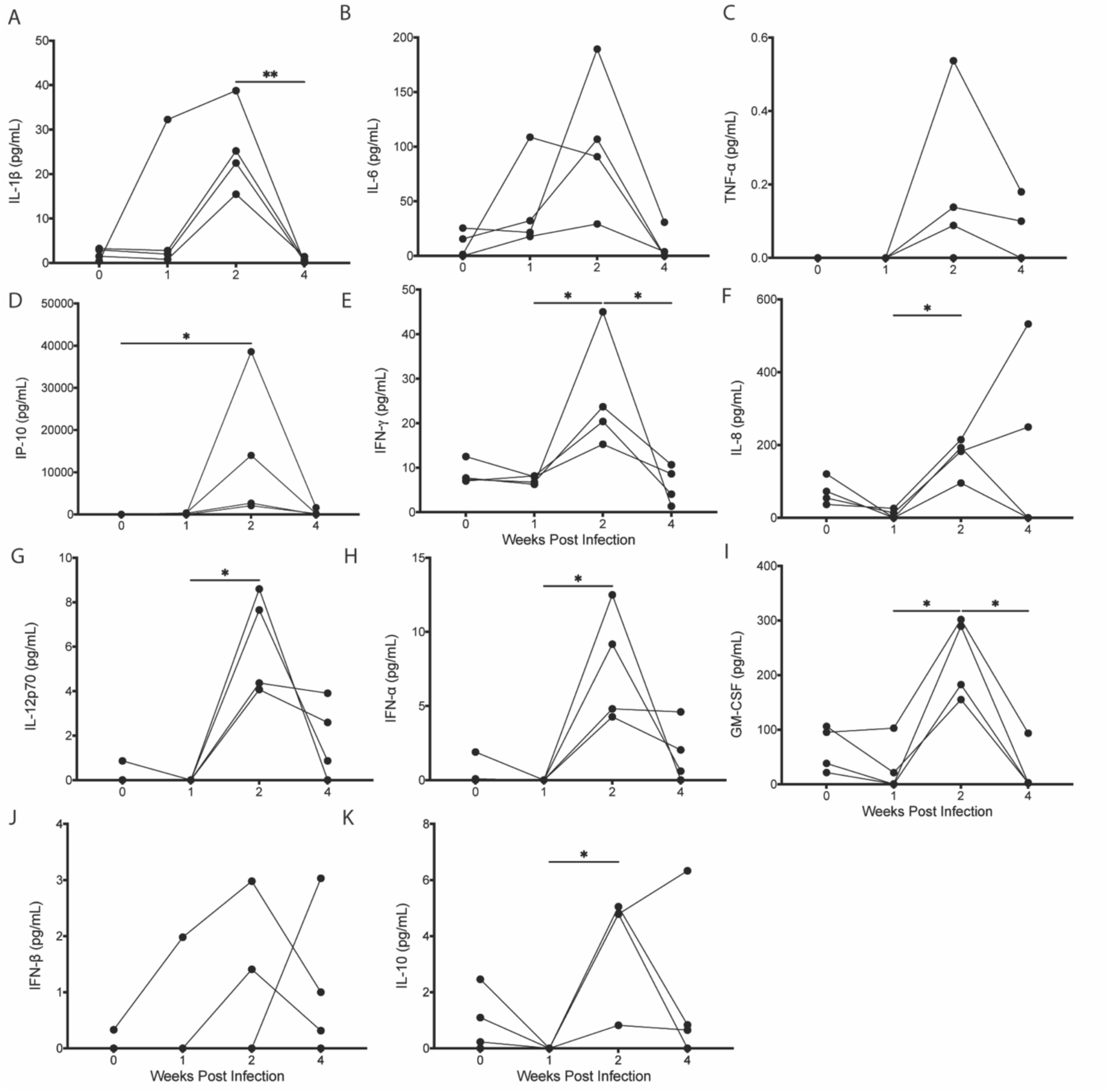
CBD Decreases Plasma Cytokines During Acute SIV Infection. Longitudinal plasma (A) IL-1β, (B) IL-6, (C) TNF-α, (D) IP-10, (E) IFN-γ, (F) IL-8, (G) IL-12p70, (H) IFN-α, (I) GM-CSF, (J) IFN-β, and (K) IL-10 determination from SIV-infected, CBD-treated macaques. *p≤0.05. **p≤0.01. Friedman test with Dunn’s multiple comparisons test. Data for each individual animal are shown.

**Supplemental Figure 4:**
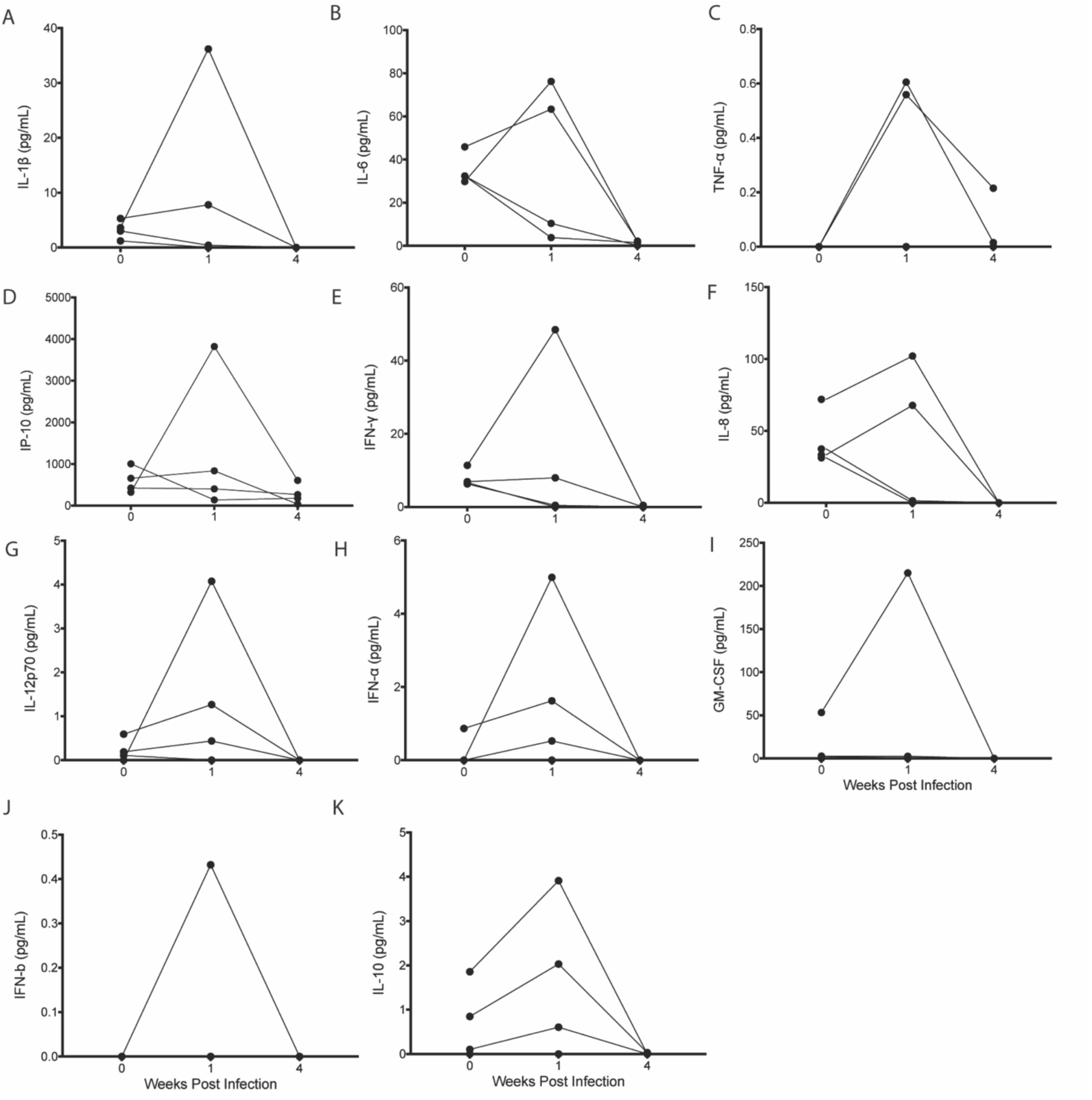
CBD Decreases CSF Cytokines During Acute SIV Infection. Longitudinal CSF (A) IL-1β, (B) IL-6, (C) TNF-α, (D) IP-10, (E) IFN-γ, (F) IL-8, (G) IL-12p70, (H) IFN-α, (I) GM-CSF, (J) IFN-β, and (K) IL-10 determination from SIV-infected, CBD-treated macaques. Data for each individual animal are shown.

**Supplemental Figure 5:**
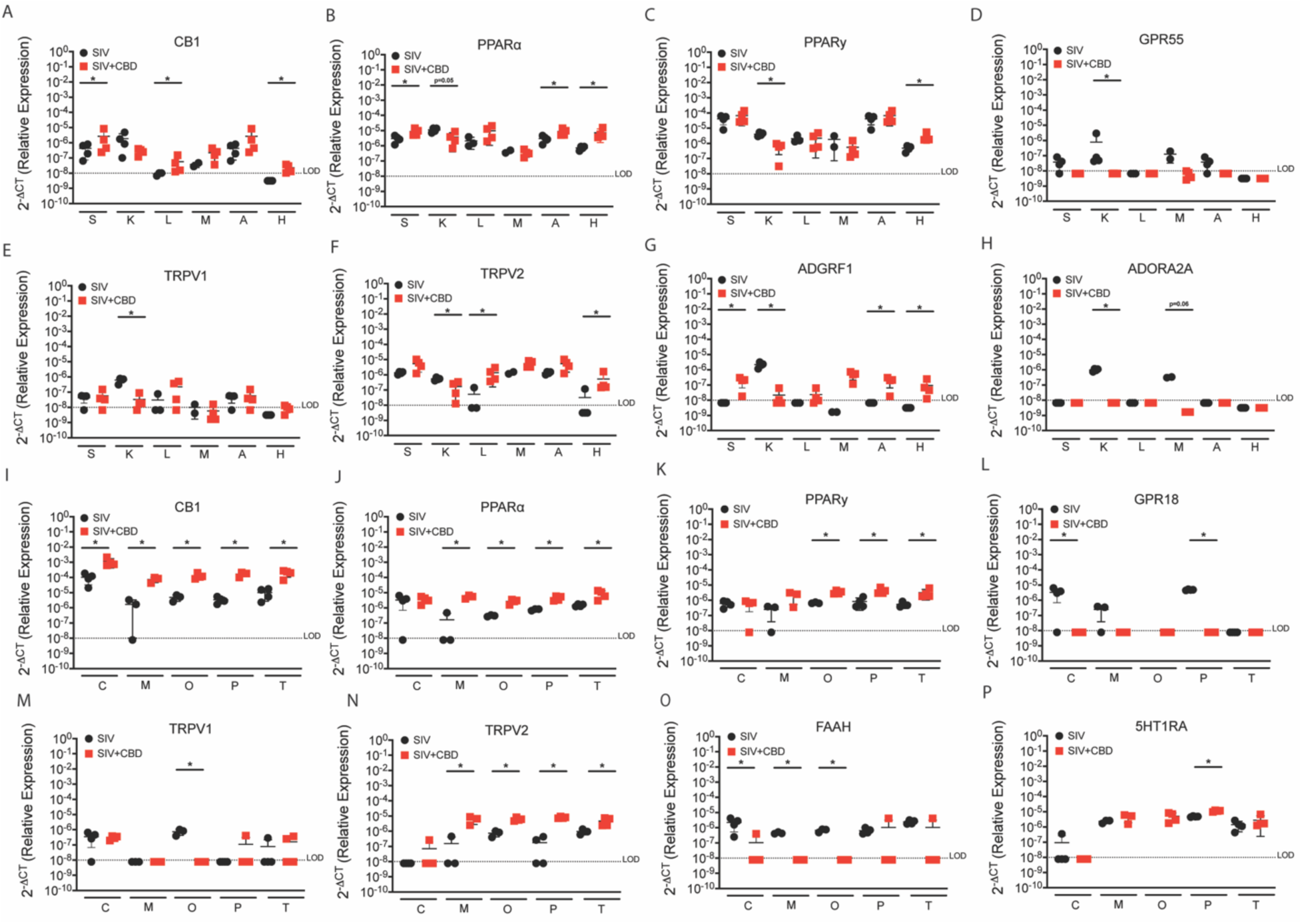
CBD Regulates Endocannabinoid Receptors During Acute SIV Infection. Relative gene expression change in (A-H) peripheral organs and (I-P) brain of (A, I) *cnr1*, (B, J) *pparα*, (C, K) *pparγ*, (D) *gpr55*, (E, M) *trpv1*, (F, N) *trpv2*, (G) *adgrf1*, (H) *adora2a*, (L) *gpr18*, (O) *faah*, and (P) *5htr1a* from SIV-infected (black) and SIV-infected, CBD-treated (red) macaques. spleen (s), kidney (k), liver (l), mesenteric lymph node (m), adipose tissue (a), heart (h), cerebellum (c), midbrain (m), occipital cortex (o), parietal cortex (p), thalamus (t). Data are represented as mean±standard deviation. *p≤0.05. Two-tailed Mann-Whitney test.

**Supplemental Figure 6:**
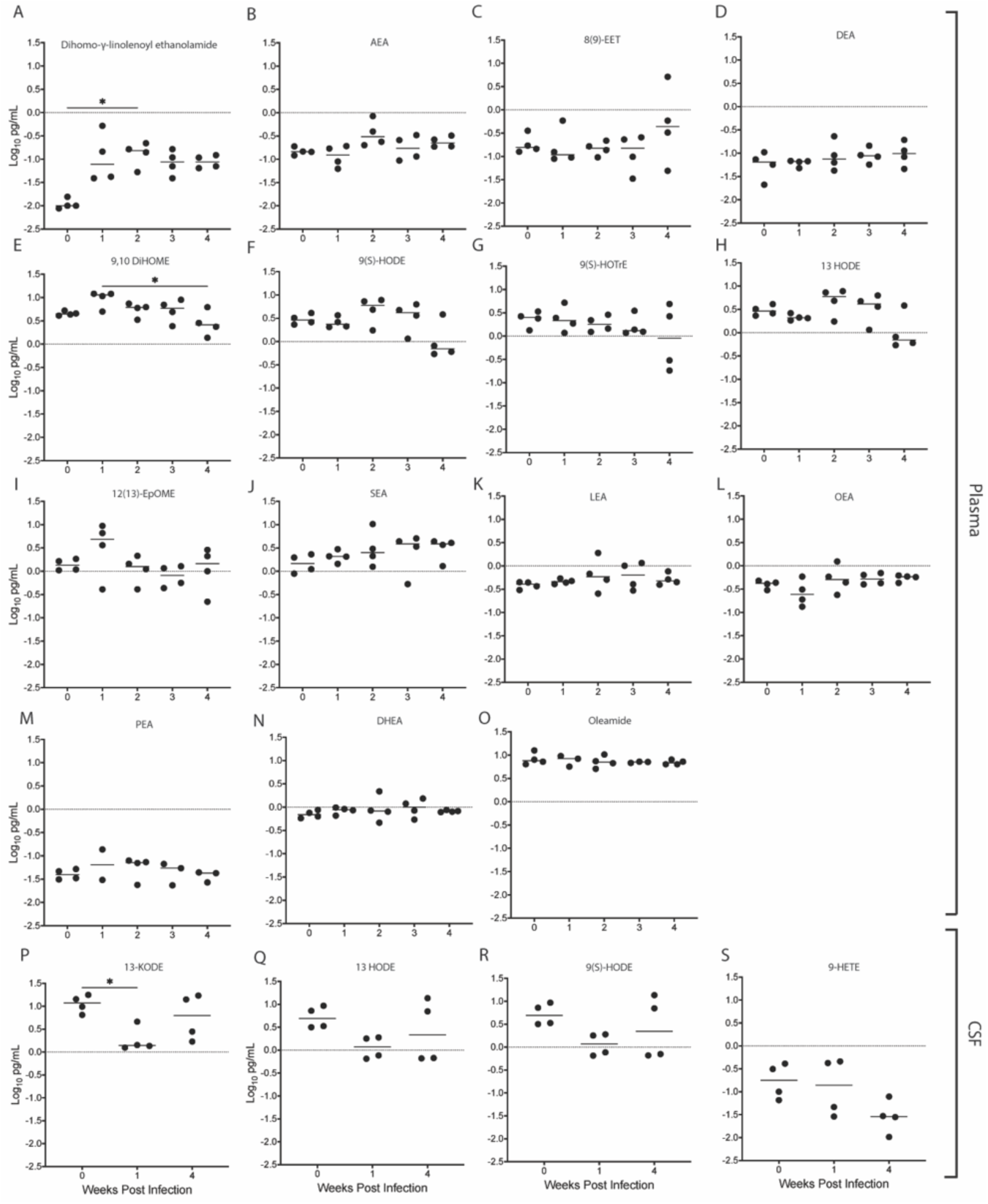
Endocannabinoid-Related Lipids Largely Remain Unchanged in SIV-Infected, CBD-Treated Macaques. Longitudinal evaluation of (A) dihomo-g-linolenoyl ethanolamide, (B) anandamide (AEA), (C) (±)8,9-epoxy-5Z,11Z,14Z-eicosatrienoic acid (8(9)-EET), (D) docosatetraenoylethanolamide (DEA), (E) (±)9,10-dihydroxy-12Z-octadecenoic acid (9,10 DIHOME), (F, R) 9- hydroxyoctadecadienoic acid (9[S]-HODE), (G) 9S-hydroxy-10E,12Z,15Z- octadecatrienoic acid (9(S)-HOTrE), (H, Q) (±)-13-hydroxy-9Z,11E-octadecadienoic acid (13 HODE), (I) (±)12(13)epoxy-9Z-octadecenoic acid (12(13)-EPOME), (J) stearoylethanolamide (SEA), (K) linoleoyl Ethanolamide (LEA), (L) oleoylethanolamide (OEA), (M) palmitoylethanolamide (PEA), (N) docosahexaenoyl ethanolamide (DHEA), (O) oleamide, and (S) (±)-9-hydroxy-5Z,7E,11Z,14Z-eicosatetraenoic acid (9-HETE) in plasma and CSF from SIV-infected, CBD-treated macaques. Data were log transformed and are represented as median values for each individual animal. *p≤0.05. Kruskal-Wallis test with Dunn’s multiple comparisons test.

**Supplemental Figure 7:**
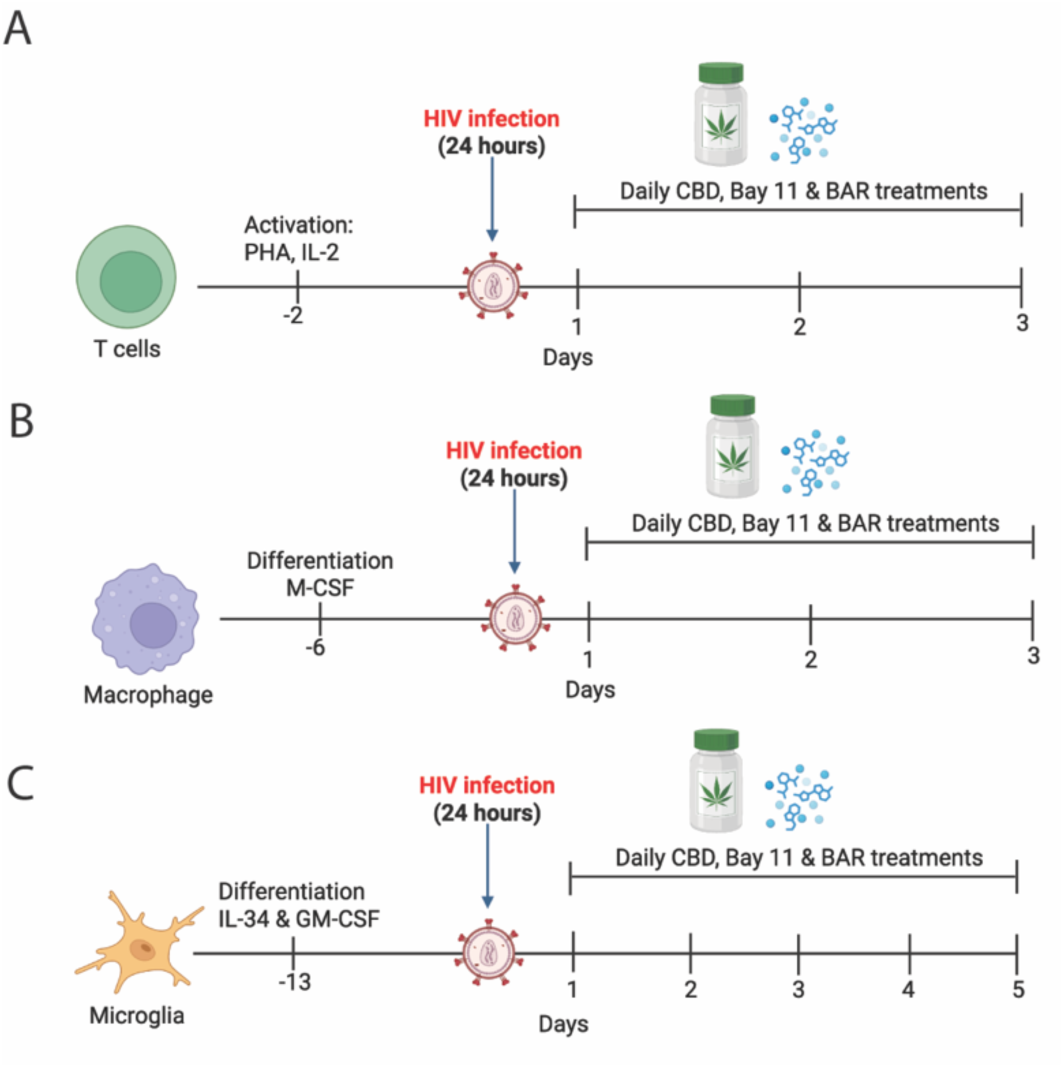
HIV Infection and CBD Treatment Paradigm for Primary Human Cells. Schematic representation of *in vitro* culture, HIV-infection, and treatment conditions for (A) T cells, (B), macrophages, and (C) microglia.

**Supplemental Figure 8:**
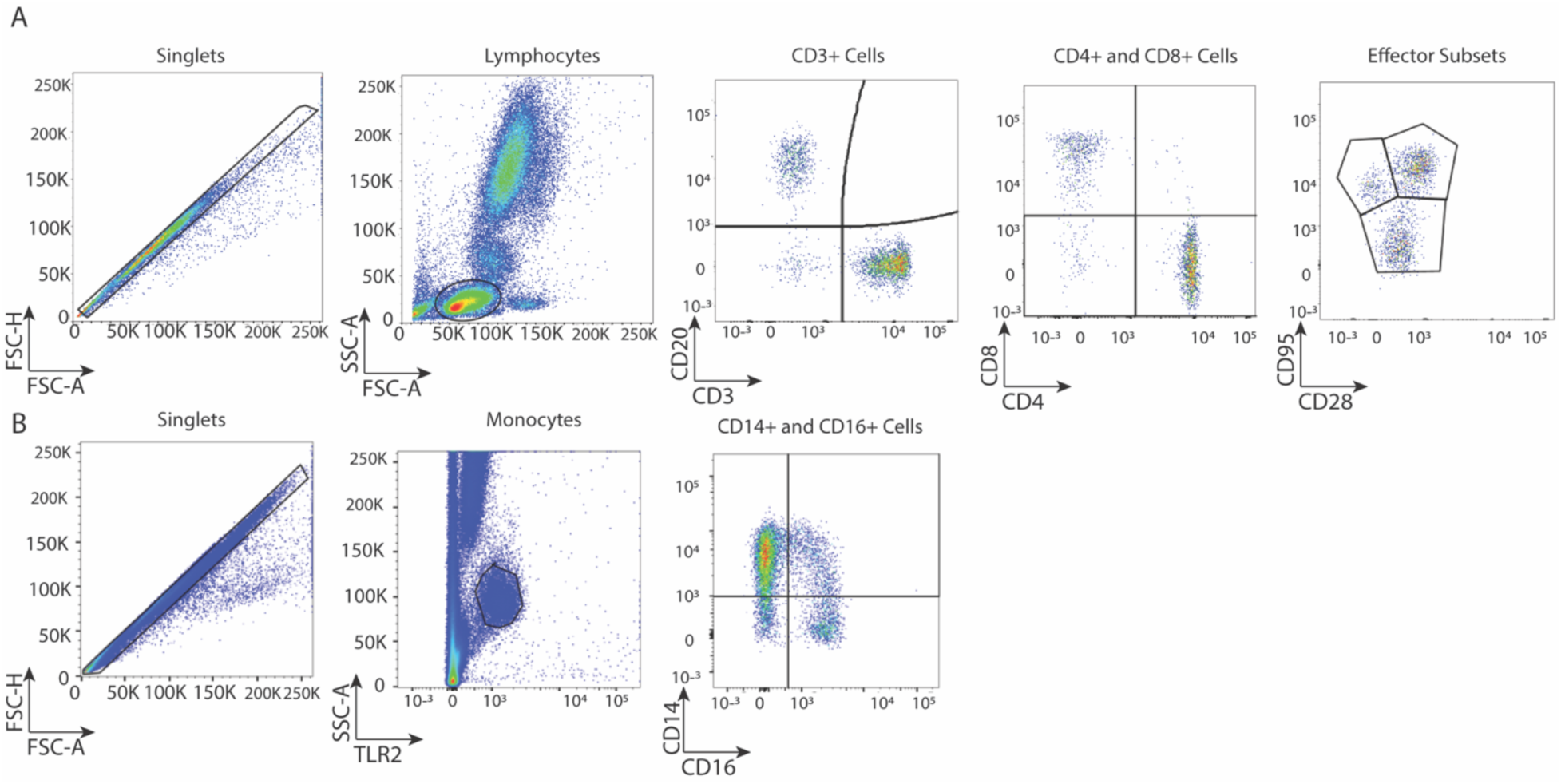
Flow Cytometry Gating Strategy of peripheral blood mononuclear cells and splenic single-cell suspensions. Evaluation of (A) lymphocyte and (B) monocyte subsets where the denoted cell populations are indicated by boxed areas. (A) Lymphocyte analysis included 1) forward scatter height (FSC-H) versus area (FSC-A) to analyze single cell events by doublet discrimination, 2) side scatter area (SSC-A) versus FSC-A to identify lymphocytes, 3) discrimination of T Cells by CD3 positivity, 4) CD3 positive cells were gated according to CD4 expression, and finally CD3+CD4+ cells were analyzed according to CD28 and CD29 positivity. (B) Monocyte analysis included 1) forward scatter height (FSC-H) versus area (FSC-A) to analyze single cell events by doublet discrimination, 2) discrimination of monocytes by TLR2 positivity, and 3) finally TLR2+ cells were analyzed according to CD14 and CD16 positivity.

